# Basis for antibody- and hormone-mediated activation of TSHR in Graves’ disease

**DOI:** 10.1101/2022.06.13.496020

**Authors:** Jia Duan, Peiyu Xu, Xiaodong Luan, Yujie Ji, Qingning Yuan, Xinheng He, Ye Jin, Xi Cheng, Hualiang Jiang, Shuyang Zhang, Yi Jiang, H. Eric Xu

## Abstract

Thyroid stimulating hormone (TSH), through activation of its G protein-coupled receptor TSHR, controls the synthesis of thyroid hormone (TH), an essential metabolic hormone. Aberrant signaling of TSHR by autoantibodies causes Graves’ disease and hypothyroidism that affect millions of patients worldwide. Here we report the active structures of TSHR with TSH and an activating autoantibody M22, both bound to an allosteric agonist ML-109, as well as an inactive TSHR structure with inhibitory antibody K1-70. Both TSH and M22 push the extracellular domain (ECD) of TSHR into the upright active conformation. In contrast, K1-70 blocks TSH binding and is incapable of pushing the ECD to the upright conformation. Comparisons of the active and inactive structures of TSHR with those of the luteinizing hormone–choriogonadotropin receptor (LHCGR) reveal a universal activation mechanism of glycoprotein hormone receptors, in which a conserved 10-residue fragment (P10) from the hinge C-terminal loop mediated interactions from the receptor ECD to its transmembrane domain. One surprisingly feature is that there are over 15 cholesterols surrounding TSHR, supporting its preferential location in lipid rafts. These structures also highlight a common mechanism for TSH and autoantibody M22 to activate TSHR, thus providing the molecular basis for Graves’ disease.

---

Thyroid stimulating hormone (TSH) is a pituitary hormone that connects the signaling cascade from the hypothalamus to the thyroid, which is termed the hypothalamic-pituitary-thyroid axis (HPT), which is critical for many physiological functions^1^ (Fig. 1a). The function of TSH is mediated through its binding and activation of the G protein-coupled receptor, TSHR, which is richly expressed in thyroid^2^. TSHR activation is primarily coupled to the Gs and Gq proteins and leads to up-regulation of the secondary message cAMP and phospholipase C^3^, which then stimulate the production and secretion of thyroid hormone, an essential hormone for metabolic homeostasis and development in vertebrates^4^ (Fig. 1a).

**Figure 1.**
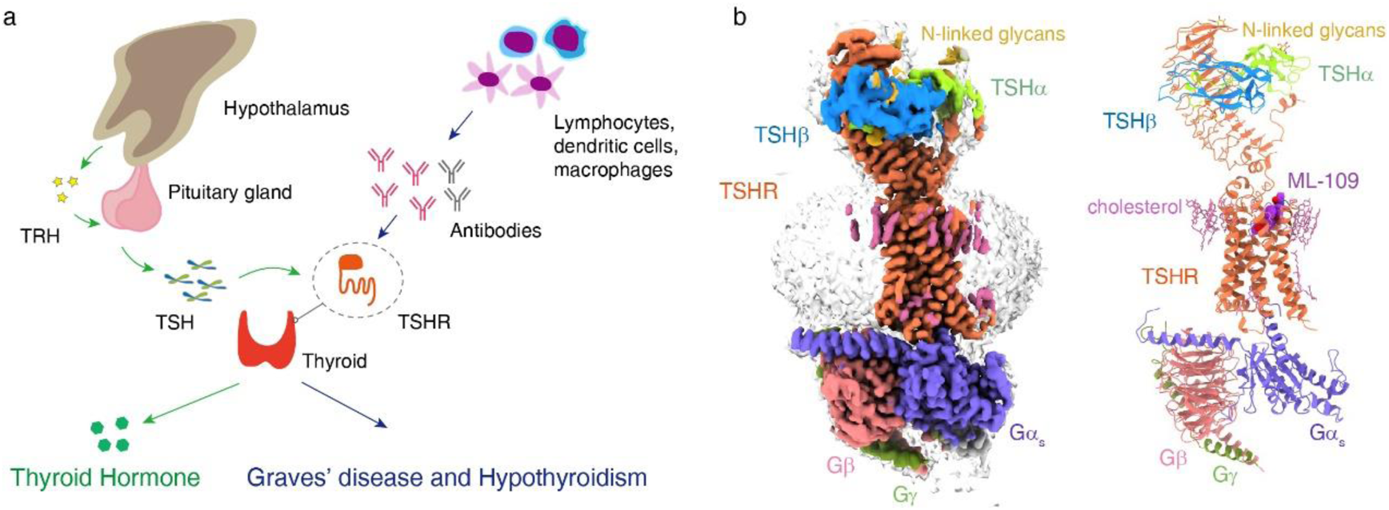
Cryo-EM structure of the TSH-TSHR-Gs complex. **a**, Schematic diagram of TSHR physiology. **b**, Cryo-EM density (left panel) and ribbon presentation (right panel) of the TSH-TSHR-Gs complex.

TSH belongs to the glycoprotein hormone family, which also includes follicle-stimulating hormone (FSH), luteinizing hormone (LH) and chorionic gonadotropin (CG)^5^. TSHR and related glycoprotein hormone receptors contain a large extracellular domain (ECD) of 11-12 leucine-rich-repeats (LRRs), a hinge region, and a transmembrane domain (TMD)^6,7^. In the case of TSHR, its ECD can be shedding from TMD and the free ECD could become an autoantigen that induces various types of antibodies against TSHR^8,9^. Autoantibodies, including M22, that activate TSHR increase abnormal production of thyroid hormone and lead to Graves’ disease^10^ (Fig. 1a). Autoantibodies such as K1-70 that inhibit TSHR decrease thyroid hormone that causes hypothyroidism and Hashimoto’s disease^10,11^ (Fig. 1a). In combination, the diseases caused by the aberrant signaling of TSHR affect over hundreds of millions of patients worldwide^12^. However, the basis of TSH-mediated TSHR activation or antibody-mediated TSHR activation and inhibition remains unknown.

Because of its disease relevance, TSHR is an attractive target of drug discovery^13^. Small molecule agonist such as ML-109, a selective TSHR allosteric agonist, has been developed^14^. In this paper, we used cryo-electron microscope (cryo-EM) to determine an extensive set of TSHR structures bound to ML-109, TSH, and autoantibodies M22 and K1-70. Combined with functional studies, our structures reveal a conserved activation mechanism of glycoprotein hormone receptors and highlight a shared mechanism for TSH and autoantibody M22 to activate TSHR, thus providing the molecular basis for Graves’ disease as well as for hypothyroidism and Hashimoto’s disease.

## Structure of the TSH-TSHR-Gs complex

To determine the mechanism of TSH-mediated TSHR activation, we first solved the structures of human TSHR bound to human TSH. TSHR used in our studies contains a deletion of residues 317-366 in the hinge region, which is naturally removed from matured TSHR^15^. It has been shown that the region of residues 317-366 is not required for TSHR folding and signaling, and its deletion increases TSHR stability^15,16^. In addition, a TSHR constitutively active mutation, S281I, was also introduced to enhance the assembly of the TSH–TSHR–Gs complex^6,17^. The structure was determined with TSH and TSHR in complex with the mini-Gs/Gβγ heterotrimer, the Gs stabilizing nanobody Nb35^18^, and the allosteric agonist ML-109, to a global nominal resolution of 2.96 Å (Fig. 1b, Extended Data Fig. 1, Table 1). Local refinement of the TSH-TSHR ECD subcomplex yielded a map at a resolution of 2.67 Å (Extended Data Fig. 1). The quality of EM map is sufficient for placement of ML-109, TSH, TSHR, the mini G heterotrimer into the structure model (Fig. 1b, Extended Data Fig. 2a). In addition, more than 20 cholesterol molecules were found to surround the TMD of the TSHR, especially near the extracellular side, like a cholesterol belt circulating the top half of the TMD (Fig 1b and Extended Data Fig. 3a). This finding is consistent with that TSHR is preferentially located within lipid rafts that are enriched with cholesterols^19,20^.

The overall structure of the TSH-TSHR complex reveals an up-right ECD conformation relative to the membrane layer (Fig. 1b), with TSH binding to the concave surface of the ECD, which contains 12 LRR units (Fig. 2a). TSH is a heterodimer of cysteine-knot protein, which shares a common α-subunit with other glycoprotein hormones and a unique β-subunit that determines hormone specificity^21–23^. The TSH structure in the complex resembles those of FSH and CG^24,25^, with an elongated fold for both α and β subunits that contains three conserved glycosylation sites (two in α-chain and one in β-chain) (Fig. 2b, Extended Data Fig. 4b,4c). The glycosylation at N52 from α-chain is at the interface between α and β subunits (Extended Data Fig. 3b), and is probably important for the stability of TSH.

**Figure 2.**
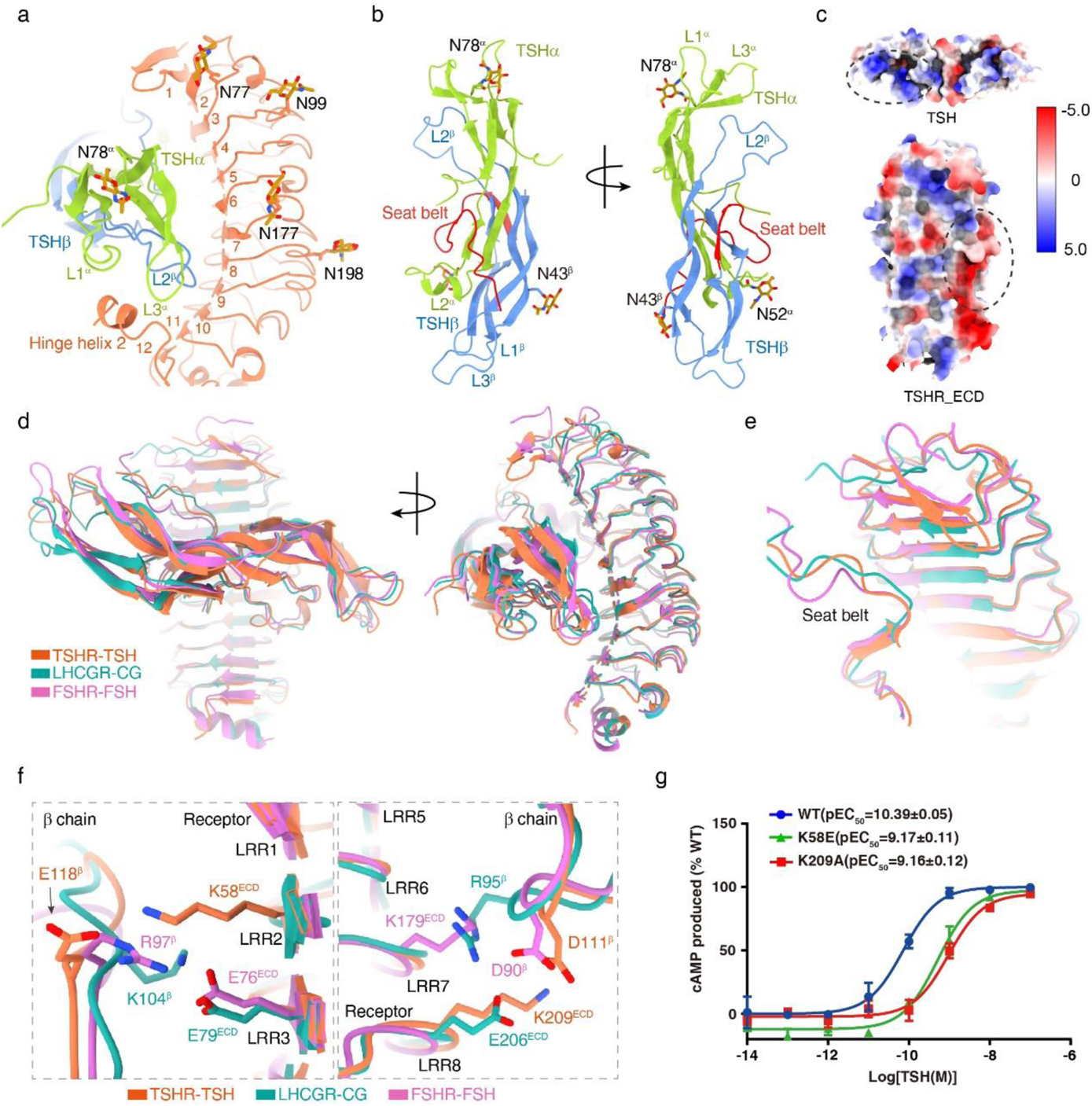
Hormone specificity and interactions between TSH and TSHR. **a,** The interface of the TSHR ECD with TSHα and TSHβ subunits. The density map of the extended C-terminus of hinge helix is shown in pink. **b**, Structure of TSHα and TSHβ subunits, the TSHβ C-terminal “seat belt” is highlighted in red and N-linked glycans are highlighted in sphere. **c**, Surface charge distribution of TSH and TSHR-ECD, the charge complementary interactions are highlighted in black circles. **d-f,** Structural comparison of the TSH-TSHR complex with the CG-LHCGR complex (PDB code: 7FIH) and FSH-FSHR complex (PDB code: 4AY9), the oval receptor ECD structures comparison in two views (**d**), seat belt comparison (**e**), detail interactions comparison (**f**). **g**, Concentration-response curves for point mutants in TSHR ECD. Data are shown as the mean ± s.e.m. from three independent measurements. WT, wild type.

## Recognition of TSH by TSHR

In the structure, TSH forms direct contact with LRR2-10 with both charge complementary interactions and shape-matching interactions (Fig. 2a, c). The binding of TSH to TSHR buries the total surface areas of 1955 Å^2^. Structure comparisons of TSH and other glycoprotein hormones bound to their receptors reveal the overall similarity in their ECD structures and the binding interfaces^6,7^ (Fig. 2d), but the major difference is seen in the C-terminal seat belt of the β-subunit (Fig. 2e). The RMSD of the Cα atoms from the TSH seat belt residues 107-121 to those of FSH seat belt residues 86-94 is 0.49 Å, with the Cα of residues 93-100 deviated from the corresponding CG by as much as 0.76 Å. The differences in the TSH seat belt structure also result in the different interaction patterns in the binding interface (Fig. 2f, Extended Data Fig. 4a, 4b), particularly D111 and E118 from the TSH seat belt, which forms salt bridges with K209 and K58 from TSHR. These electrostatic interactions are specific to the TSH-TSHR complex and mutations in K209 and K58 of TSHR resulted in a more than a 10-fold reduction of TSH potency (Fig. 2g, Extended Data Table 2), supporting their important role in TSH binding specificity and receptor activation.

At the C-terminus of the TSHR ECD is the hinge region, which contains LRR11, two α-helices (termed as hinge helix 1 and 2), LRR12, a short linker fragment (residues 396-404), and the conserved p10 region (residues 405-414) (Extended Data Fig. 3c, 4b). Together with LRR1-10, the hinge region forms the complete TSHR ECD. Following the conserved p10 region is the TSHR TMD, which adopts the active conformation and is coupled to the G protein heterotrimer at its cytoplasmic face. The TSH-TSHR structure is highly similar to the active structure of the CG-LHCGR complex^6^ (Fig. 3a, Extended Data Fig. 3d), with the RMSD of their TMD (residues 415-697) less than 0.53 Å, suggesting that both TSH and CG stabilize the receptor active conformation through a common mechanism, including stabilizing the conformation of the conserved p10 region^26^. Interestingly, the TSHR ECD is rotated by additional ~12° to the upright position relative to the ECD of the active LHCGR structure (Fig. 3a).

**Figure 3.**
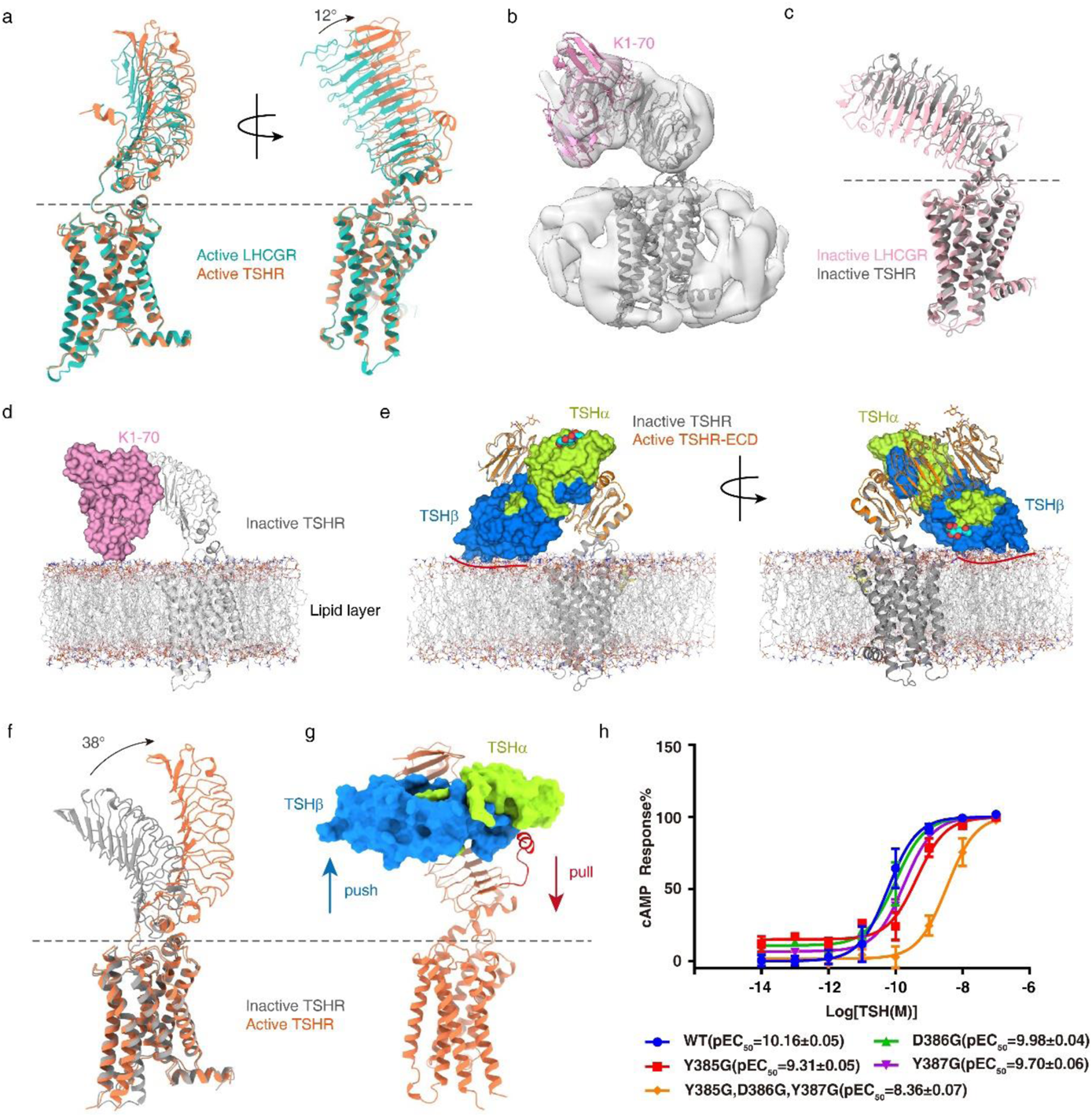
Basis for TSH-mediated TSHR activation. **a,** Structural comparison of the active full-length TSHR and LHCGR, the ECD rotation angle was measured at the Cα atom of residue V87 and I281 from TSHR and D84 from LHCGR. **b,** EM density and ribbon presentation of K1-70_ScFv-TSHR. **c**, Comparison of inactive full-length TSHR and LHCGR, the ECD rotation angle was measured at the Cα atom of residue V87 and S281 from TSHR and K109 from LHCGR. **d**, The putative model of K1-70_ScFv-TSHR in the membrane layer, K1-70_ScFv is shown in surface. **e**, The putative model of TSH interacts with inactive TSHR in the membrane layer, TSH is shown in surface and two N-linked glycans are shown in sphere, the clash of TSHβ with the membrane layer is marked with a red line. **f**, Structural comparison of the active and inactive TSHR, the ECD rotation angle was measured at the Cα atom of residue N74 and S281 from inactive TSHR and N74 from active TSHR. **g**, The Schematic diagram of TSH-induced TSHR activation. **h**, Concentration-response curves for point mutants in hinge helix 2. Data are shown as the mean ± s.e.m. from three independent measurements. WT, wild type.

## TSH-mediated TSHR activation

To fully understand TSH-induced TSHR activation, we attempted to solve the inactive structure of TSHR. Through numerous trials, we were only able to obtain a 5.46 Å structure of TSHR in the presence of a blocking antibody K1-70, isolated from a patient with hypothyroidism^27^, and a small molecule antagonist Org 274179-0^28^ (Extended Data Fig. 5). The relatively low resolution of the density map prevented accurate modeling of detailed structures of TSHR and Org 274179-0, but is sufficient for us to place the ECD, the TMD as well as the ScFv fragment of K1-70 (Fig. 3b), which was able to block TSHR activation by TSH^8,29^. The ScFv of K1-70 appeared to bind to the distal N-terminal region of the TSHR ECD (LRR1-7) as reported by the previous X-ray crystal structure^29^, with overlapping areas with the TSH binding interface (Extended Data Fig. 3e), thus blocking TSH binding (Extended Data Fig. 3f). Structure comparison of the TSHR-K1-70 complex with the inactive LHCGR structure reveals that the ECD of TSHR has a ~8.6° rotation away from the membrane layer relative to the ECD of the inactive LHCGR (Fig. 3c). In this position, the ScFv fragment of K1-70 does not clash with the membrane layer (Fig. 3d). In contrast, superposition of the ECD of the active TSHR to that of the inactive TSHR reveals that the L3^β^ loop of TSH will still clash with the membrane layer (Fig. 3e), in an analogous manner to CG binding to LHCGR, despite that the TSHR ECD is rotated ~8.6° from the membrane layer. Structure comparison of the active and inactive TSHR reveals that TSH binding induces a ~38° rotation of the ECD from the inactive position to the active upright conformation (Fig. 3f). These results thus highlight the common mechanism for glycoprotein hormones to activate their receptors by pushing the ECD to rotate away from the membrane layer to the upright active conformation.

In addition to the push model of TSHR activation, its ECD is also pulled by the hinge region, in which residues 382-390 adopt a short α-helix (Fig. 2a, Extended Data Fig. 1d) to interact with TSH in a manner similar to what has been proposed for FSH-FSHR^7^ and CG-LHCGR^6^. The region of 382-390 contains a negative residue D386 and two tyrosine residues, which could be sulfonated^7^. The negative charge property of this region has been proposed to interact with the positive charge region of R54 from the TSH beta chain (Extended Data Fig. 3g). Individual mutations of D386, Y385 and Y387 result in a 5-10 fold reduction of TSH activation potency, and an 80-fold reduction in combined mutated receptors (Fig. 3h, Extended Data Table 2), supporting the common pull mechanism for glycoprotein hormone to activate their receptors. Together, TSH-mediated TSHR activation shares a similar “push and pull” model that has been proposed for CG-induced LHCGR activation (Fig. 3g). Interestingly, shortening the hinge loop by deleting 104 loop residues (291-394) resulted in a constitutively active receptor (Extended Data Fig. 3h), suggesting that the flexible hinge loop could prevent the ECD from adopting the active conformation in the apo receptor.

## Autoantibody-mediated TSHR activation

To uncover the mechanism of autoantibody-induced TSHR activation, we solved the structures of TSHR bound to the small molecule allosteric agonist ML-109 and an ScFv fragment of M22, a human monoclonal autoantibody to the TSHR from a patient with Graves’ disease^30^. This ScFv also exhibited potent activity to activate TSHR, with similar efficacy and potency of TSH (Fig. 4a). The structure was determined to a global resolution of 2.78 Å, and local refinement of the M22_ScFv-TSHR ECD subcomplex yielded a map at a resolution of 2.39 Å (Extended Data Fig. 6). The clear density map that allows the placement of M22_ScFv and cholesterol molecules into the TSHR-Gs complex (Fig. 4b, Extended Data Fig. 2b, 2c, Table 1). The M22_ScFv-TSHR binding interface is similar to the previous structure of M22 binding to a TSHR ECD fragment (residues 1-260) that contain LRR1-10^31^. In the full-length TSHR complex, the M22_ScFv-TSHR binding interface is extended to the LRR11-12 region that is part of the hinge region near the TMD interface (Fig. 4c). The total buried surface area by the M22_ScFv binding is 1419 Å^2^. Superposition of the TSHR ECD from the M22_ScFv bound structure with the inactive ECD structure reveals that the heavy chain of M22_ScFv will clash with the membrane layer (Fig 4d), consistent with the push model of the TSHR activation. However, interaction with the hinge regions was not observed, thus antibody-mediated TSHR activation is not mediated through a pull mechanism as seen by activation with the endogenous hormone such as TSH and CG. It is likely that M22_ScFv binding has a larger interface with TSHR, thus pulling by the hinge region is not required for M22-mediated activation. Indeed, mutations in the hinge region (D386G, Y385G and Y387G) affect TSH activation but not antibody-mediated activation (Fig. 4e, Extended Data Table 2).

**Figure 4.**
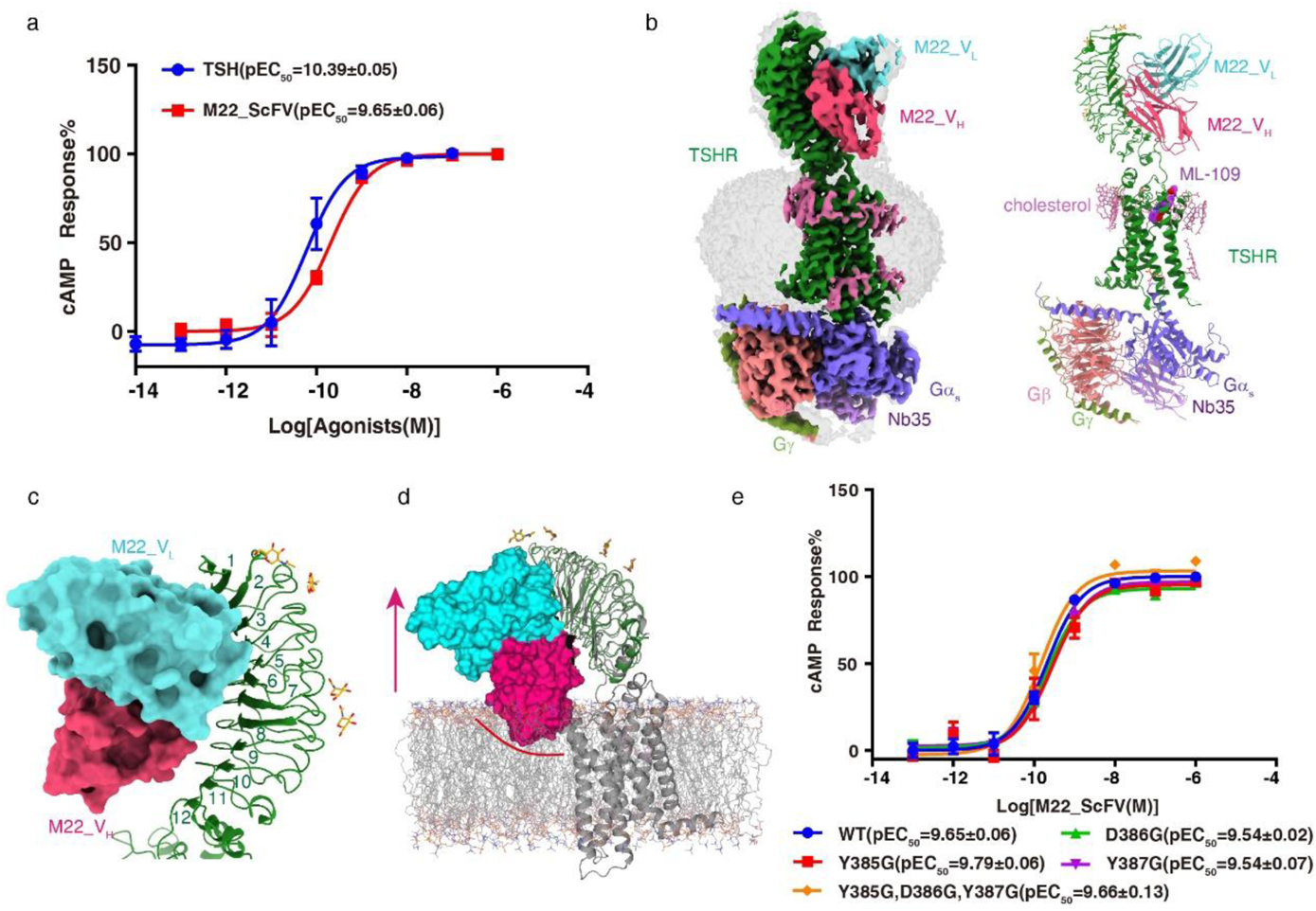
Basis for M22_ScFv mediated TSHR activation. **a**, Concentration-response curves for TSH and M22_ScFv induced TSHR activation. Data are shown as the mean ± s.e.m. from three independent measurements. WT, wild type. **b**, Cryo-EM density (left panel) and ribbon presentation (right panel) of the M22_ScFv-TSHR-Gs complex. **c**, The interface of the M22_ScFv with TSHR ECD. M22_ScFv is shown in surface. **d**, The putative model of M22_ScFv interacts with inactive TSHR in the membrane layer, M22_ScFv is shown in surface and the clash of the heavy chain with the membrane layer is marked with a red line. **e**, Concentration-response curves for point mutants in hinge helix 2. Data are shown as the mean ± s.e.m. from three independent measurements. WT, wild type.

## TSHR activation by an allosteric agonist

In both TSH- and M22-bound TSHR structures, the small molecule allosteric agonist ML-109 was added to stabilize the complex. ML-109 is known to stimulate thyroid function in human thyrocytes and mice^14^. Consistently, ML-109 activated TSHR with full efficacy (Extended Data Fig. 7a) and was able to synergistically act with TSH and M22 on TSHR (Extended Data Fig. 7b, 7c). The density map for the ML-109 in the M22_ScFv-bound complex is clear for defining the binding mode (Fig. 5a) and is thus used for our presentation below. ML-109 binds to a large pocket formed by TM helices 3-7 within the top half of the TMD (Fig. 5b, 5c). In addition, the long ECL2 together with ECL3 and the P10 loop form the open entry of the pocket for ML-109 to dock into (Fig. 5c). The interactions of ML-109 with TSHR are almost exclusively hydrophobic (Fig. 5d, 5e), with the phenyl-acetamide sitting at the top entry of the pocket to interact with M572 from ECL2. The central phenyl-methoxy group is packed between M572 from ECL2 on one side and I640 from TM6 on the other side. The quinazolinyl group binds to the bottom of the pocket with its hydroxyl pointed to V586 from TM5. The methyl benzyl group interacts with I648 from TM6 and M572 from ECL2. Consistently, mutations at these residues resulted in the reduction of the activation potency of ML-109 to TSHR (Fig. 5f, Extended Data Fig. 7d, 7e, Table 2).

**Figure 5.**
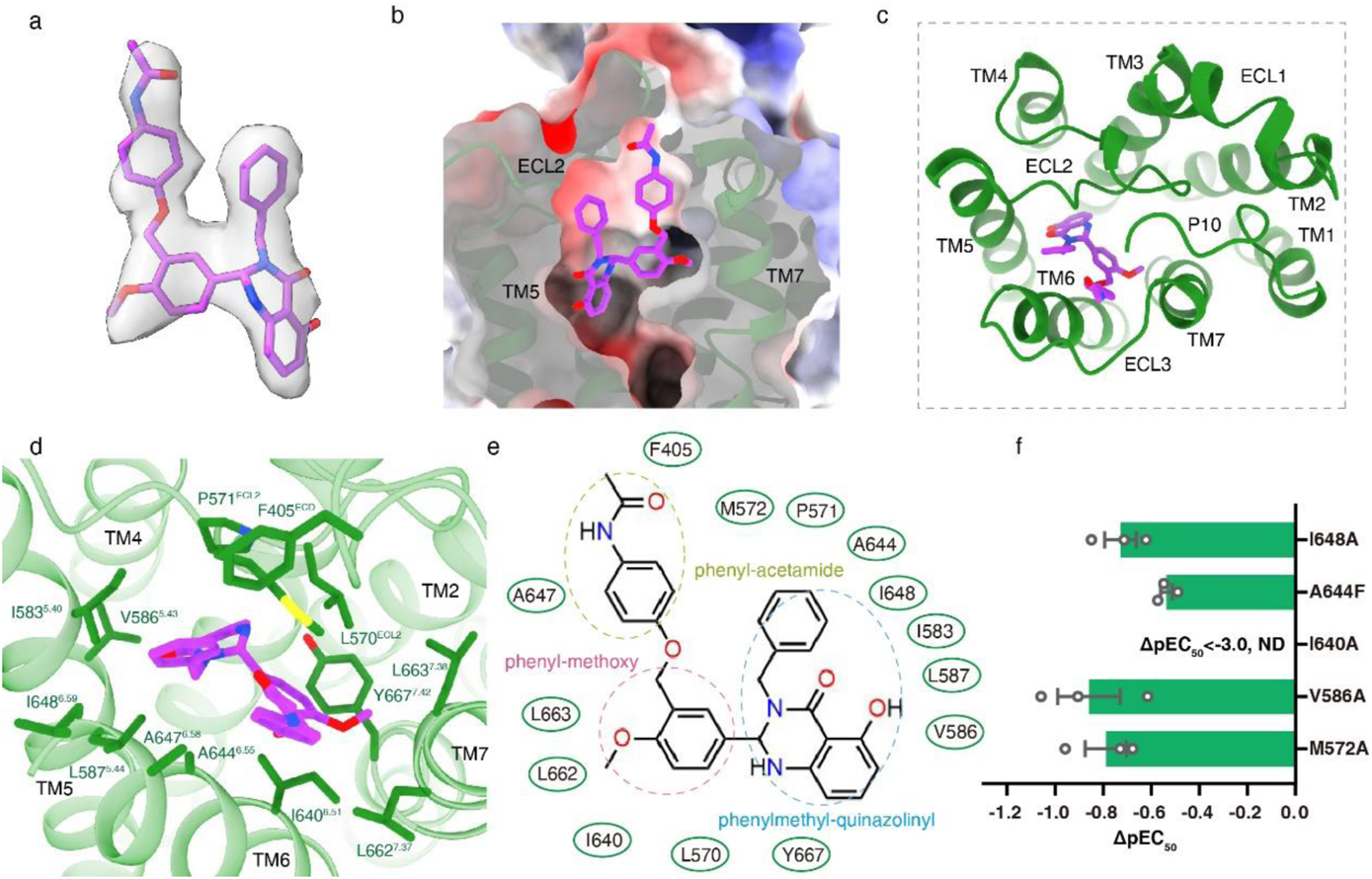
Basis for TSHR activation by ML-109. **a,** The EM density of ML-109. **b-c,** The binding pocket of ML-109 in TSHR, from the front view (**b**) and top view (**c**). **d**, Detail interactions between ML-109 and TSHR. **e**, Structure of ML-109 and schematic representation of ML-109-TSHR interactions. **f**, Effects of different pocket mutations on the efficacy of ML-109-induced cAMP accumulation. Data are shown as the ΔpEC_50_ ± s.e.m. from three independent measurements.

ML-109 is a TSHR-specific allosteric agonist that is not active for LHCGR and FSHR^14^. Structure comparison of ML-109 bound TSHR with Org43553 bound LHCGR reveals very similar pocket and ligand binding modes between these two structures (Extended Data Fig. 7f). The major difference is a 2.2 Å inward shift of LHCGR at the residue A593 from the C-terminal end of TM6 relative to that of TSHR (Extended Data Fig. 7f). This inward shift has caused steric collisions of A589 and A592 from TM6 of LHCGR with the central phenyl-methoxy group and the bottom quinazolinyl group of ML109, thus excluding ML-109 from binding to LHCGR.

## Discussion

In this paper, we reported three structures of TSHR bound to its natural hormone TSH, synthetic allosteric agonist ML-109, autoimmune disease antibodies M22 and K1-70 from patients with Graves’ disease and hypothyroidism^27,30^, respectively. Analyses of these structures reveal that TSH binding to TSHR induces a 38° rotation of its ECD toward the upright active position, analogous to a 45° rotation seen in the CG-LHCGR structure. Similarly, the upright active conformation of the TSH-TSHR structure is further stabilized by interactions of the bound TSH with a hinge helix from the receptor. Together, these observations highlight the universal mechanism of “push and pull” for activation of glycoprotein hormone receptors.

TSHR is a unique GPCR with a large ECD that has numerous autoantibodies from patients with Graves’ disease and hypothyroidism. The structures of TSHR bound with M22 and K1-70 provide the first glimpse into the mechanism for activation and inhibition of a G protein-coupled receptor by autoantibodies. Based on the structure observations, M22-mediated TSHR activation appears to act through three combined mechanisms: 1) by destabilizing the inactive conformation as the M22 binding mode to the inactive TSHR clashes with the membrane layer, thus its binding would induce rotation of its ECD to the upright position; 2) by binding site close to the ECD-TMD interface, thus stabilizing the upright active conformation; and 3) by enlarging the binding interface from LRR2 to LRR12, thus alleviating the requirement of the pull from the hinge loop region. In contrast, the blocking antibody K1-70 binds to the far distal N-terminal LRR 1-7, which overlaps TSH binding but is compatible with the inactive ECD conformation that is similar to the inactive LHCGR structure, thus providing the mechanistic explanation for the blocking activity of K1-70. Together, these structures provide a basis for antibody-mediated activation and inhibition of TSHR, which is highly relevant to human diseases associated with the imbalance of thyroid hormones.

Our structures also reveal the binding mode of the TSHR small molecule allosteric agonist ML-109, which is docked into the TMD pocket that is mostly conserved in the glycoprotein hormone receptors, with a similar binding mode to Org43553 seen in the LHCGR structure. However, the ML-109 binding pocket in TSHR is distinct from the Org43553 binding pocket in LHCGR, particularly the inward shift of the C-terminal end of TM6, which causes clashes of LHCGR with ML-109, therefore providing the basis for the selectivity of ML-109 for TSHR.

One of the most intriguing aspects of TSHR is its tendency to be prone to active and inactive mutations that are highly associated with hyperthyroidism and hypothyroidism^10,32^. The locations of these active mutations are spread across in the TSHR TMD, with rich distribution within the TM6 region (Extended Data Fig. 8), suggesting that the dynamic of TM6 is key to TSHR activation. In agree with this notion, mutations of I640A and A644F above also resulted in high levels of constitutive activation. Both of these mutations are from TM6 and modeling of these two mutations suggests that they affect the packing and stability of TM6 (Extended Data Fig. 7e, Table 2), thus highlighting the important role of TM6 in the receptor activation.

In summary, the structures of TSHR with endogenous hormone TSH and small molecule agonist ML-109 reveal the mechanism of hormone and agonist induced receptor activation. The structures with the activating antibody M22 and the blocking antibody K1-70 also uncover the basis of how TSHR can be up- or down-regulated by autoantibodies, thus providing a basis for antibody drug discovery targeting this therapeutically important receptor. In addition, the large numbers of cholesterols surrounding the TSHR TMD provide a mechanistic explanation for the long-historic observations that TSHR is preferentially located in cholesterol-rich lipid rafts for its signaling.

## Method

### Constructs

Human TSHR (full-length TSHR with residues 21-764, except residues 317-366 were removed) was cloned with an N-terminal FLAG and C-terminal His8 tags using homologous recombination (CloneExpress One Step Cloning Kit, Vazyme). Additional mutation S281I was designed to form TSHR-Gs complexes. The native signal peptide was replaced with the haemagglutinin (HA) to increase protein expression. The M22 and K1-70 antibodies were modified into ScFv epitopes, and cloned into pFastBac with a GP67 signal peptide and C-terminal His8 tag. A dominant-negative bovine Gαs construct was generated based on mini-Gs^33^. Additionally, three mutations (G226A, A366S and L272D) were also incorporated by site-directed mutagenesis to decrease the affinity of nucleotide-binding and increase the stability of Gαβγ complex^34^. All the three G-protein components, including rat Gβ1 and bovine Gγ2, were cloned into a pFastBac vector, respectively.

### Expression and purification of Nb35

Nanobody-35 (Nb35) with a C-terminal His6 tag, was expressed and purified as previously described^18^. Nb35 was purified by nickel affinity chromatography (Ni Smart Beads 6FF, SMART Lifesciences), followed by size-exclusion chromatography using a HiLoad 16/600 Superdex 75 column and finally spin concentrated to 5 mg/ml.

### Expression and purification of M22_ScFv and K1-70_ScFv

Purification of M22_ScFv and K1-70_ScFv were conducted similar as ScFv16 as previously described^35^ with a subtle change. Briefly, secreted M22_ScFv and K1-70_ScFv from baculovirus-infected Hi5 insect cells (Invitrogen) were purified using nickel affinity and size exclusion chromatography. The Ni-NTA eluted samples were collected and applied to a HiLoad Superdex 200, 10/60 column (GE Healthcare). The monomeric peak fractions were concentrated to 8 mg/mL and fast-frozen by liquid nitrogen.

### Complex expression and purification

TSHR, Gαs, Gβ1 and Gγ2 were co-expressed in *Sf9* insect cells (Invitrogen) using the Bac-to-Bac baculovirus expression system (ThermoFisher). Cell pellets were thawed and lysed in 20 mM HEPES, pH 7.4, 100 mM NaCl, 10% glycerol, 5 mM MgCl_2_ and 5 mM CaCl_2_ supplemented with Protease Inhibitor Cocktail, EDTA-Free (TargetMol). The TSH-TSHR-Gs complex was formed in membranes by the addition of 0.5 μM TSH (Thyrogen, the TSH is a recombinant protein expressed from CHO cells with natural human sequence, and is a product used in clinics.), 10 μM ML-109 (TargetMol), 10 μg/mL Nb35 and 25 mU/mL apyrase. The suspension was incubated for 1.5 h at room temperature. The membrane was then solubilized using 0.5% (w/v) n-dodecyl β-D-maltoside (DDM, Anatrace), 0.1% (w/v) cholesterol hemisuccinate (CHS, Anatrace) and 0.1%(w/v) sodium cholate for 2 h at 4 °C. The supernatant was collected by centrifugation at 80,000 × g for 40 min and then incubated with M1 anti-Flag affinity resin for 2 h at 4 °C. After batch binding, the resin was loaded into a plastic gravity flow column and washed with 10 column volumes of 20 mM HEPES, pH 7.4, 100 mM NaCl, 10% glycerol, 2 mM MgCl_2_, 2 mM CaCl_2_, 0.01% (w/v) DDM, 0.002%(w/v) CHS, and 0.002%(w/v) sodium cholate, 0.05 μM TSH and 5 μM ML-109, further washed with 10 column volumes of same buffer plus 0.1%(w/v) digitonin, and finally eluted using 0.2 mg/mL Flag peptide. The complex was then concentrated using an Amicon Ultra Centrifugal Filter (MWCO 100 kDa) and injected onto a Superdex200 10/300 GL column (GE Healthcare) equilibrated in the buffer containing 20 mM HEPES, pH 7.4, 100 mM NaCl, 2 mM MgCl_2_, 2 mM CaCl_2_, 0.05 (w/v) digitonin, 0.0005% (w/v) sodium cholate, 0.01 μM TSH and 1 μM ML-109. For M22_cFv bound complex, 1 μM M22_ScFv was added for complex formation and 0.1 μM M22_ScFv for the following procedures. The complex fractions were collected and concentrated for electron microscopy experiments, respectively.

### K1-70_ScFv-TSHR complex expression and purification

*Sf9* insect cells were infected with TSHR baculovirus and cultured for 48 h at 27 °C before collection. Cell pellets were thawed and lysed in 20 mM HEPES, pH 7.4, 100 mM NaCl, 10% glycerol, supplemented with Protease Inhibitor Cocktail, EDTA-Free. The purification procedures were similar to the M22_ScFv-TSHR-Gs complex, a small molecular antagonist Org 274179-0 was also in the purification buffer with 10 μM concentration.

### cAMP response assay

The full-length TSHR (21–764) and mutants were cloned into pcDNA6.0 vector (Invitrogen) with a FLAG tag at its N-terminus. CHO-K1 cells (ATCC, #CCL-61) were cultured in Ham’s F-12 Nutrient Mix (Gibco) supplemented with 10% (w/v) fetal bovine serum. Cells were maintained at 37 °C in a 5% CO_2_ incubator with 200,000 cells per well in a 12-well plate. Cells were grown overnight and then transfected with 1 μg TSHR constructs by FuGENE® HD transfection reagent in each well for 24 h. cAMP accumulation was measured using the LANCE cAMP kit (PerkinElmer) according to the manufacturer’s instructions. The transfected cells were seeded onto 384-well plates with 2500 cells each well, and then incubated with ligands for 30 min at 37 °C, then Eu and Ulight were added separately before cAMP levels were measured. Fluorescence signals were measured at 620 and 665 nm by an Envision multilabel plate reader (PerkinElmer). Datas were analyzed using Graphpad Prism7.0, the three-parameter, nonlinear regression equation in Prism suite was used in fitting. Experiments were performed at least three times, the detail information were attached in the figure legends, each experiment conduceted in triplicate. Datas were presented as means ± SEM.

### Detection of surface expression of TSHR mutants

The cell seeding and transfection followed the same method as the cAMP response assay. After 24 h of transfection, cells were washed once with PBS and then detached with 0.2% (w/v) EDTA in PBS. Cells were blocked with PBS containing 5% (w/v) BSA for 15 min at room temperature before incubating with primary anti-Flag antibody (diluted with PBS containing 5% BSA at a ratio of 1:150, Sigma) for 1 h at room temperature. Cells were then washed three times with PBS containing 1% (w/v) BSA and then incubated with anti-mouse Alexa-488-conjugated secondary antibody (diluted at a ratio of 1:1,000, Invitrogen) at 4 °C in the dark for 1 h. After another three times of washing, cells were collected, and fluorescence intensity was quantified in a BD Accuri C6 flow cytometer system (BD Biosciences) through a BD Accuri C6 software1.0.264.21 at excitation 488 nm and emission 519 nm. Approximately 10,000 cellular events per sample were collected and data were normalized to the wild type TSHR. Experiments were performed at least three times, datas were presented as means ± SEM.

### Cryo-EM grid preparation and data collection

For the preparation of cryo-EM grids, 3 μL of the purified protein at 20 mg/mL for the TSH-TSHR-Gs complex, 30 mg/mL for the M22_ScFv-TSHR-Gs complex and 20 mg/mL for the K1-70-TSHR complex, were applied onto a glow-discharged holey carbon grid (Quantifoil R1.2/1.3). Grids were plunge-frozen in liquid ethane using Vitrobot Mark IV (Thermo Fischer Scientific). Frozen grids were transferred to liquid nitrogen and stored for data acquisition.

Cryo-EM imaging of the TSH-TSHR-Gs complex was performed on a Titan Krios at 300 kV in Cryo-Electron Microscopy Research Center, Shanghai Institute of Materia Medica, Chinese Academy of Sciences (Shanghai China), and cryo-EM imaging of the M22_ScFv-TSHR-Gs complex and K1-70-TSHR complexes were performed on a Titan Krios at 300 kV in the Advanced Center for Electron Microscopy at Shanghai Institute of Materia Medica, Chinese Academy of Sciences (Shanghai China).

A total of 14,965 movies for the TSH-TSHR-Gs complex was collected with a Gatan K3 Summit direct electron detector with a Gatan energy filter (operated with a slit width of 20 eV) (GIF) at a pixel size of 1.071 Å using the SerialEM software^36^. The micrographs were recorded in counting mode at a dose rate of about 22 e/Å^2^/s with a defocus ranging from −1.2 to −2.2 μm. The total exposure time was 3 s and intermediate frames were recorded in 0.083 s intervals, resulting in a total of 36 frames per micrograph.

A total of 6,911 movies for the M22_ScFv-TSHR-Gs complex and 5,938 movies for the K1-70-TSHR, were collected by a Gatan K3 Summit direct electron detector with a Gatan energy filter (operated with a slit width of 20 eV) (GIF) at a pixel size of 0.824 Å and 1.04 Å using the EPU software. The micrographs were recorded in counting mode with a defocus ranging from −1.2 to −2.2 μm. The total exposure time was 3.33 s and intermediate frames were recorded in 0.104 s intervals, resulting in a total of 36 frames per micrograph.

### Image processing and map construction

Dose-fractionated image stacks were aligned using MotionCor2.1^37^. Contrast transfer function (CTF) parameters for each micrograph were estimated by Gctf^38^. For the TSH-TSHR-Gs complex, particle selections for 2D and 3D classifications were performed using RELION-3.1^39^. Automated particle picking yielded 15,576,249 particles that were subjected to 3 rounds reference-free 2D classification to discard poorly defined particles, producing 2,396,330 particles. The map of the LHCGR-Gs-Nb35-CG-Org43553 complex (EMDB-31597) was used as initial reference for 3D classification, resulting in two subsets with ECD and TSH. The selected subsets were performed 3 rounds 3D classification to remove particles without clear ECD. The well-defined subsets were subsequently subjected to 3D refinement and Bayesian polishing. The final particles were performed non-uniform refinement in CryoSPARC^40^ and generated a map with an indicated global resolution of 3.04 Å with 751,617 particle projections at a Fourier shell correlation of 0.143. DeepEMhancer was used for generated a sharpen map to observation on the TSH and TSHR-hinge region.

For the M22_ScFv-TSHR-Gs complex, the cryo-EM data analysis was performed in CryoSPARC^40^. Automated particle picking yielded 2,699,981 particles that were subjected to reference-free 2D classification to discard poorly defined particles, producing 931,674 particles. The well-defined subsets were further subjected to 2 rounds hetero-refinement and a 3D classification for remove the particles without ECD. The particles were subsequently subjected to non-uniform refinement. The global refinement map shows an indicated resolution of 2.96 Å at a Fourier shell correlation of 0.143.

For the K1-70-TSHR complex, automated particle picking yielded 6,988,946 particles that were subjected to 4 rounds reference-free 2D classification to discard poorly defined particles. The LHCGR map (EMDB-31599) low-pass filtered to 60 Å was used as the initial reference for hetero-refinement, resulting in two well-defined subsets. The selected subsets were performed additional 3D classification, resulting a well-defined subset, which were subsequently subjected to non-uniform refinement. The final refinement generated a map with an indicated global resolution of 5.46 Å with 52,479 particle projections at a Fourier shell correlation of 0.143.

### Model building and refinement

For the TSH-TSHR-Gs and M22_ScFv-TSHR-Gs complexes, the structure of LHCGR-Gs-Nb35-CG-Org43553 (PDB code: 7FIH), the structure of TSHR-ECD-M22 (PDB code: 3G04) were used as the start for model rebuilding and refinement against the electron microscopy map. For the K1-70_ScFv-TSHR complex, the AlphaFold model of TSHR and the structure of TSHR-ECD-K1-70 (PDB code: 2XWT) were used as the start for model rebuilding and refinement against the electron microscopy map. The model was docked into the electron microscopy density map using Chimera^41^, followed by iterative manual adjustment and rebuilding in COOT^42^ and ISOLDE^43^. Real space and reciprocal space refinements were performed using Phenix programs^44^. The model statistics were validated using MolProbity^45^. The final refinement statistics were validated using the module “comprehensive validation (cryo-EM)” in Phenix. The final refinement statistics are provided in Extended Data Table 1. Structural figures were prepared in ChimeraX^46^ and PyMOL (https://pymol.org/2/).

### Construction of membraned TSHR

TSHR was firstly oriented by the Orientations of Proteins in Membranes server^47^. Then, the structure was inserted in 100 Å * 100 Å 1-palmitoyl-2-oleoyl-sn-glycero-3-phosphocholine (POPC) membrane according to CHARMM-GUI server^48^.

## Acknowledgments

The cryo-EM data were collected at the Cryo-Electron Microscopy Research Center and Advanced Center for Electron Microscopy, Shanghai Institute of Materia Medica (SIMM). The authors thank the staff at the SIMM Cryo-Electron Microscopy Research Center and Advanced Center for Electron Microscopy for their technical support. This work was partially supported by Ministry of Science and Technology (China) grants (2018YFA0507002 to H.E.X.); Shanghai Municipal Science and Technology Major Project (2019SHZDZX02 to H.E.X.); Shanghai Municipal Science and Technology Major Project (H.E.X.); CAS Strategic Priority Research Program (XDB37030103 to H.E.X.); the National Natural Science Foundation of China (32130022 to H.E.X., 32171187 to Y.J., 82121005 to H.E.X. and Y.J.); CAMS Innovation Fund for Medical Sciences (2021-I2M-1-003 to S.Z.); CAMS Innovation Fund for Medical Sciences (2021-CAMS-JZ004 to S.Z.); Tsinghua University-Peking University Center for Life Sciences (045-61020100121 to S. Z.); National Science & Technology Major Project “Key New Drug Creation and Manufacturing Program” of China (2018ZX09711002 to H.J.); Science and Technology Commission of Shanghai Municipal (20431900100 to H.J.); Jack Ma Foundation (2020-CMKYGG-05 to H.J.).

## Author contributions

J.D. designed the expression constructs, purified the TSHR proteins, prepared the final samples for negative stain, performed cryo-EM grid preparation and data collection, conducted functional studies, and participated in map calaulations, figure and manuscript preparation; P.X. performed cryo-EM data calculations, model building, and participated in figure preparation; X.L. helped conceived the project, supplied the TSH hormone and Org 274179-0; Q.Y. participated in cryo-EM data calculations, Y-J.J participated in functional studies, X.H. participated in figure preparation; H.J. and X.C. supervised X.H. in figure preparation; Y.J. supervised the studies, and participated in manuscript preparation; S.Z. helped conceived the project, and supervised X.L. and Y. Jin; H.E.X. conceived and supervised the project, analyzed the structures, and wrote the manuscript with inputs from all authors.

## Data availability

The density maps and structure coordinates have been deposited to the Electron Microscopy Database (EMDB) and the Protein Data Bank (PDB) with accession number of EMD-33491, PDB ID 7XW5 for the TSH-TSHR-Gs complex; EMD-33492 and 7XW6 for the M22-TSHR-Gs complex; EMD-33493 and 7XW7 for the K1-70-TSHR complex.

---

**Extended Data Fig. 1.**
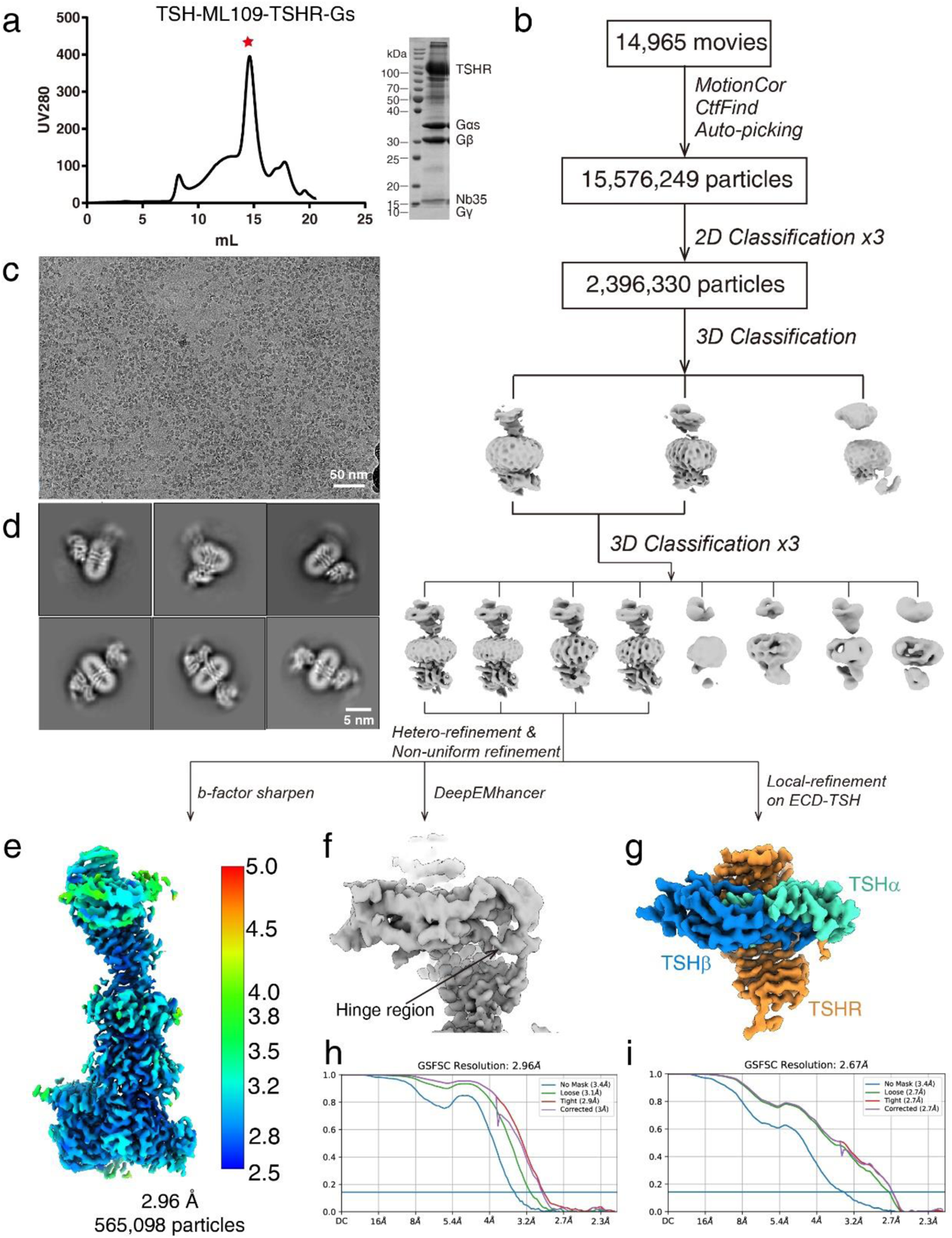
Cryo-EM images and single-particle reconstruction of the TSH-TSHR-Gs complex. **a**, Size-exclusion chromatography elution profile and SDS-PAGE of the TSH-TSHR-Gs complex. Red star indicates the monomer peak of the complex. **b-d,** Cryo-EM micrograph, reference-free 2D class averages, and flowchart of cryo-EM data analysis of the TSH-TSHR-Gs complex. **e,** Cryo-EM map of the TSH-TSHR-Gs complex colored by local resolutions from 2.5 Å (blue) to 5.5 Å (red). **f**, The density map of TSH-TSHR ECD subcomplex from DeepEMhancer analysis. **g**, The density map of TSH-TSHR ECD subcomplex from local refinement. **h, i,** The “Gold-standard” Fourier shell correlation (FSC) curve indicates that the overall resolution of the electron density map of the TSH-TSHR-Gs complex is 2.96 Å, and the local resolution of the electron density map of the TSH-TSHR ECD subcomplex is 2.67 Å.

**Extended Data Fig. 2.**
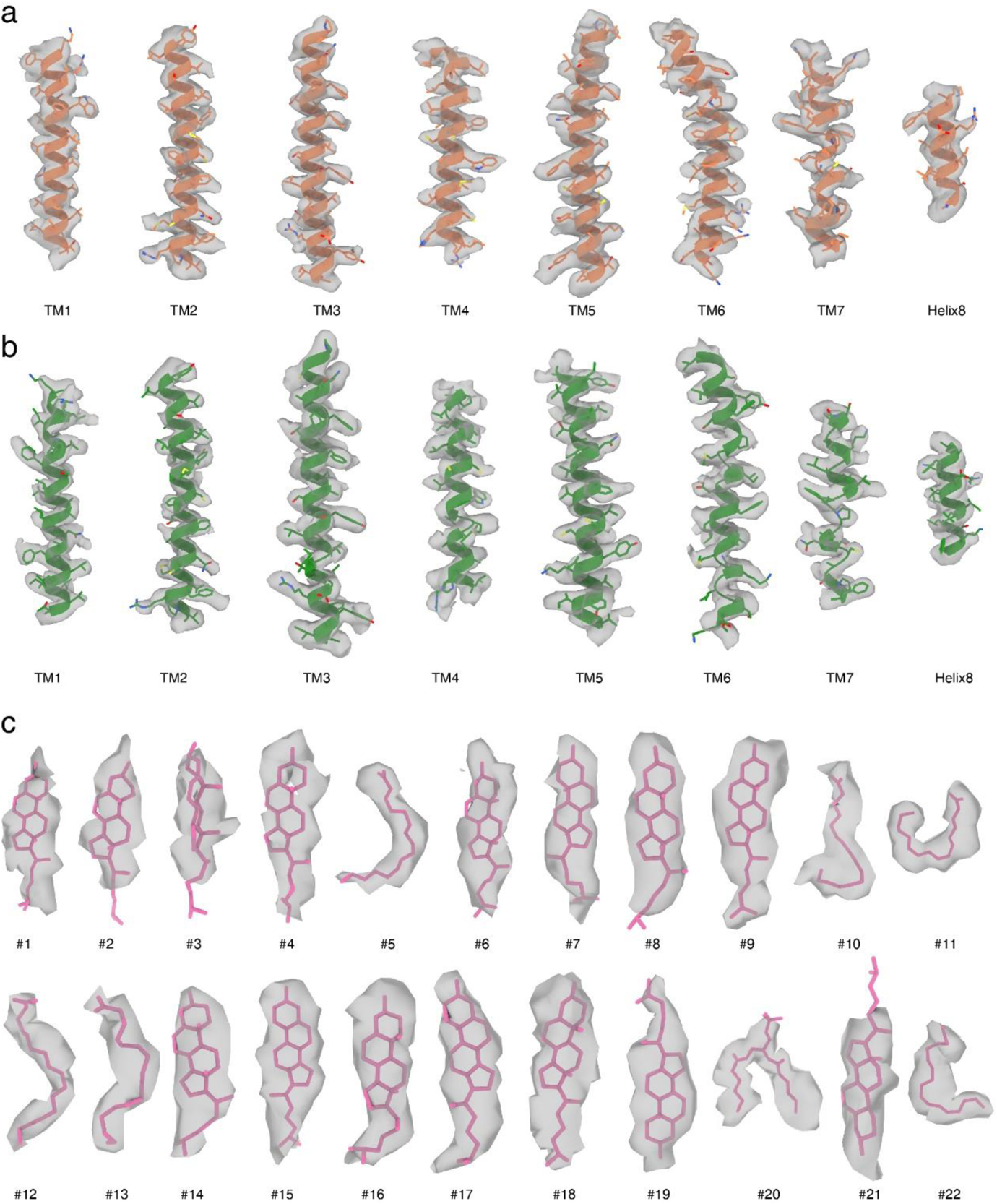
Cryo-EM image density maps with all transmembrane helices, and H8. **a**, TSHR TMD density maps in TSH-TSHR-Gs complex; **b**, TSHR TMD density maps in M22_ScFv-TSHR-Gs complex. **c**, Cholesterol and lipid density maps in M22_ScFv-TSHR-Gs complex.

**Extended Data Fig. 3.**
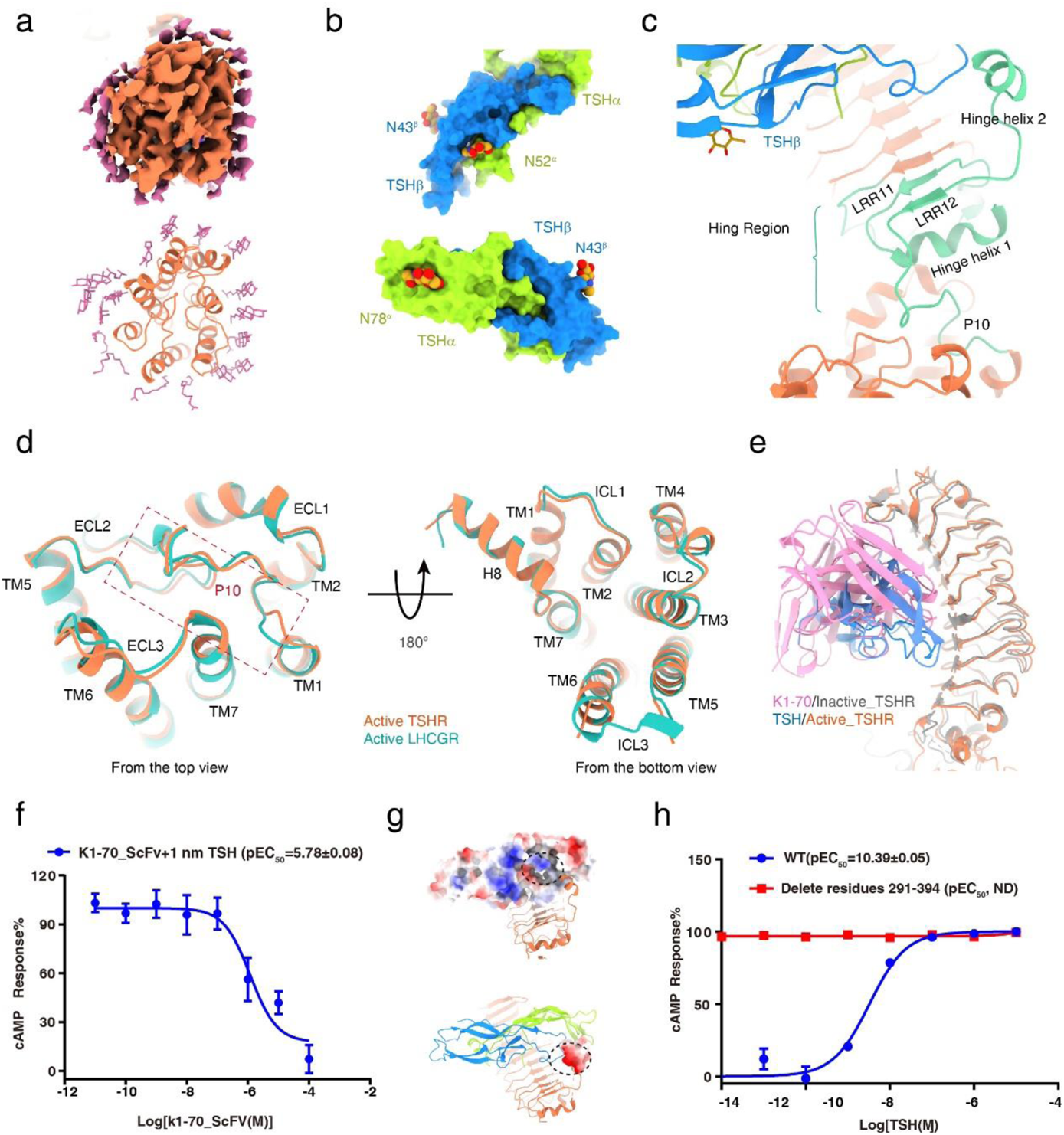
Structural features of TSHR in TSH-TSHR-Gs complex. **a**, Cryo-EM density (top panel) and ribbon presentation (bottom panel) of cholesterol molecules around the TSHR TMD in the TSH-TSHR-Gs complex. **b**, Surface representation of TSHα and TSHβ subunits, the three N-linked glycans are shown in sphere. **c**, Ribbon presentation of hinge region in TSHR. **d**, Structure comparison of active TSHR and active LHCGR TMD, top view (**c**) and bottom view (**d**). **e,** Structure alignment of TSH-TSHR ECD and K1-70-TSHR ECD. The binding interface of TSH overlaps with K1-70_ScFv. **f**, Concentration-response curves for TSHR cAMP accumulation with K1-70_ScFv and 1 nm TSH. **g**, The positively charged pockets in TSH and negatively charged hinge helix 1 surface are highlighted in black circles. **h**, Concentration-response curves for TSHR hinge loop deletion. Data from three independent experiments are presented as the mean ± s.e.m.

**Extended Data Fig. 4.**
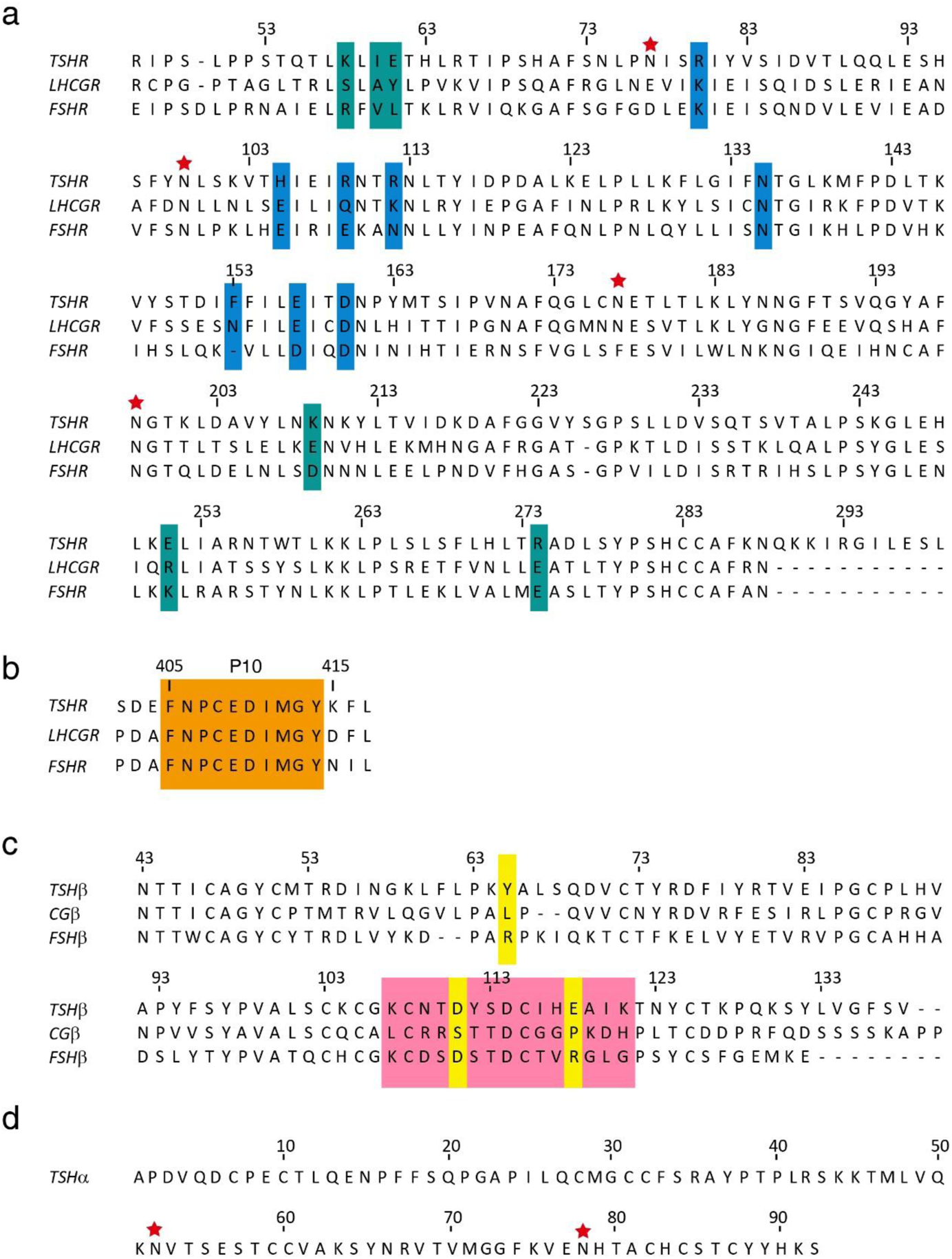
Sequence alignment of glycoprotein hormones and related receptors. **a**, Sequences alignment of human TSHR, LHCGR and FSHR in the region of the hormone-binding domain. Residues interact with TSH labeled in light sea green and blue, residues that determine TSH-TSHR specificity were labeled in light sea green. **b**, Sequences alignment of human TSHR, LHCGR and FSHR in the region of the P10 fragments. P10 is shown in green. **c**, Sequences alignment of human TSH, CG and FSH β subunit. The may interface of TSHβ interacted with TSHR are highlighted in red, residues that determine TSH-TSHR specificity were labeled in yellow. **d**, The α-subunit sequence of glycoprotein hormones. Structure resolved N-linked glycans are highlighted with red stars.

**Extended Data Fig. 5.**
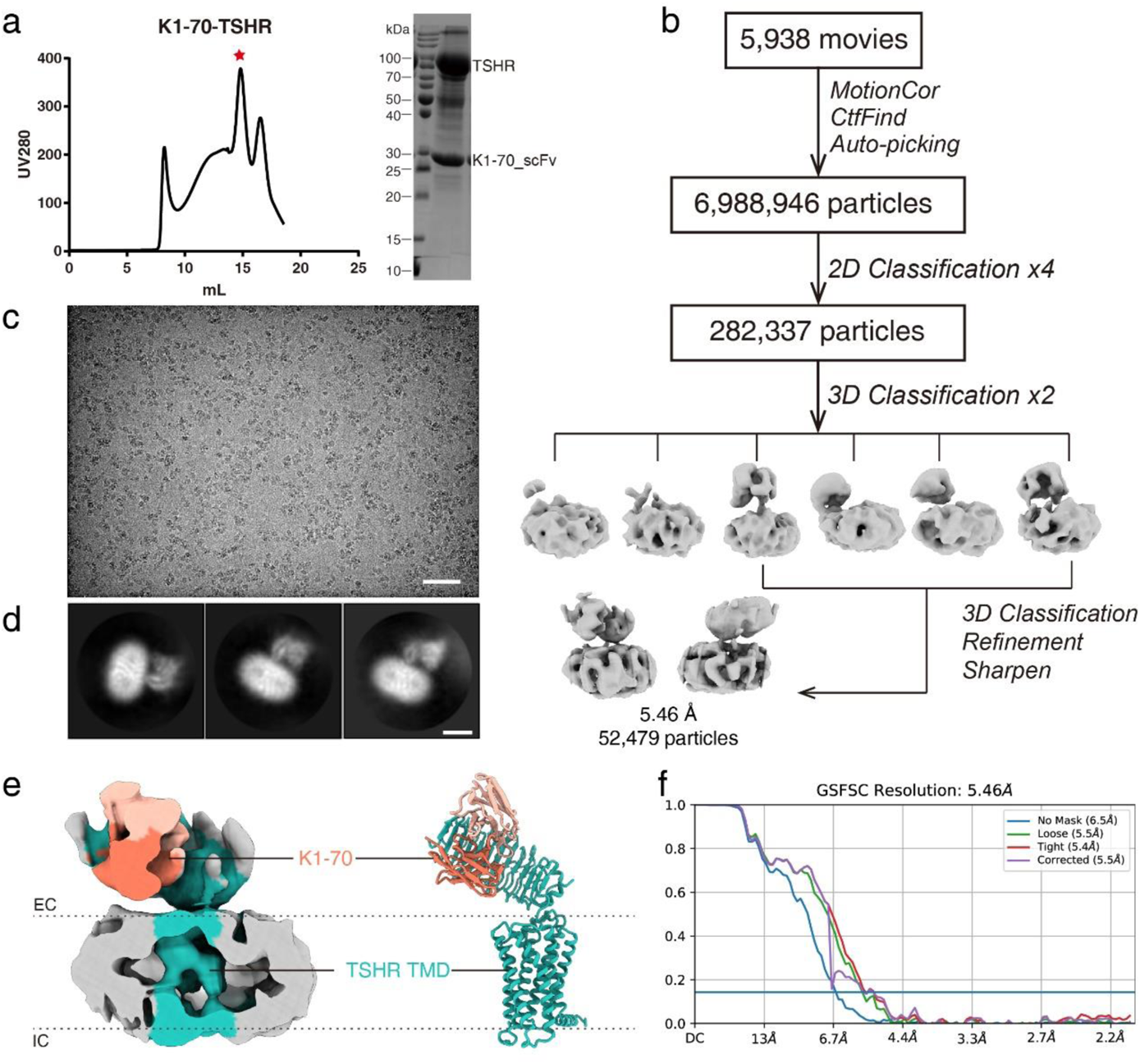
Cryo-EM image and single-particle reconstruction of the K1-70_ScFv-TSHR complex. **a**, Size-exclusion chromatography elution profile and SDS-PAGE of the K1-70_ScFv-TSHR complex. Red star indicates the monomer peak of the two complex. **b-d,** Cryo-EM micrograph, reference-free 2D class averages, and flowchart of cryo-EM data analysis of the K1-70_ScFv-TSHR complex. **e**, K1-70_ScFv-TSHR complex map and model. **f**, The “Gold-standard” Fourier shell correlation (FSC) curves indicate that the overall resolution of the K1-70_ScFv-TSHR complex is 5.46 Å.

**Extended Data Fig. 6.**
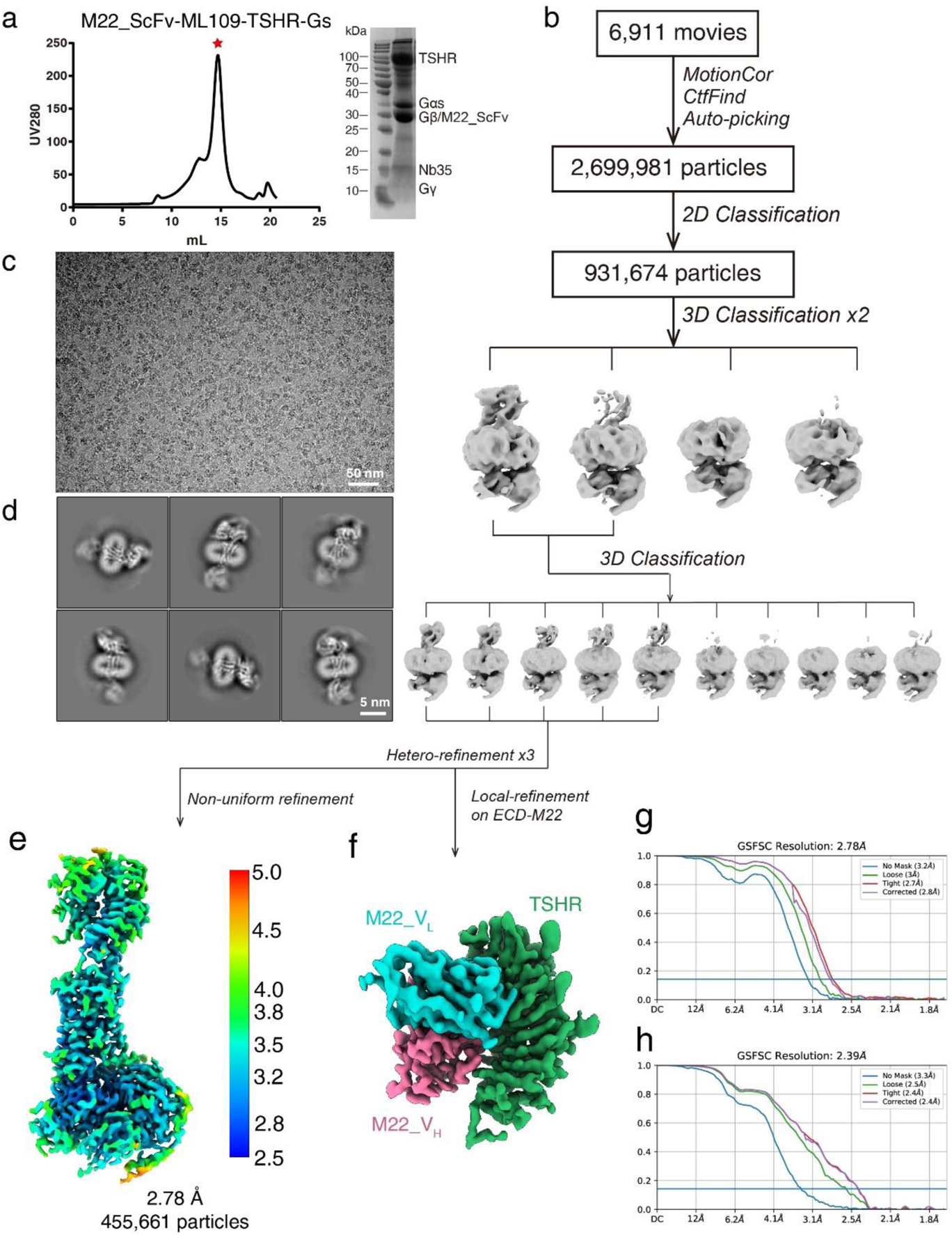
Cryo-EM images and single-particle reconstruction of the M22_ScFv-TSHR-Gs complex. **a**, Size-exclusion chromatography elution profiles and SDS-PAGEs of the M22_ScFv-TSHR-Gs complex. Red star indicates the monomer peak of the complex. **b-d,** Cryo-EM micrograph, reference-free 2D class averages, and flowchart of cryo-EM data analysis of the M22_ScFv-TSHR-Gs. **e,** Cryo-EM map of the M22_ScFv-TSHR-Gs complex colored by local resolutions from 2.5 Å (blue) to 5.0 Å (red). **f**, Cryo-EM map of the M22_ScFv-TSHR ECD subcomplex from local refinement. **g, h,** The “Gold-standard” Fourier shell correlation (FSC) curves indicate that the overall resolution of the electron density map of the M22_ScFv-TSHR-Gs complex is 2.78 Å, and the local resolution of the electron density map of the M22_ScFv-TSHR ECD subcomplex is 2.39 Å.

**Extended Data Fig. 7.**
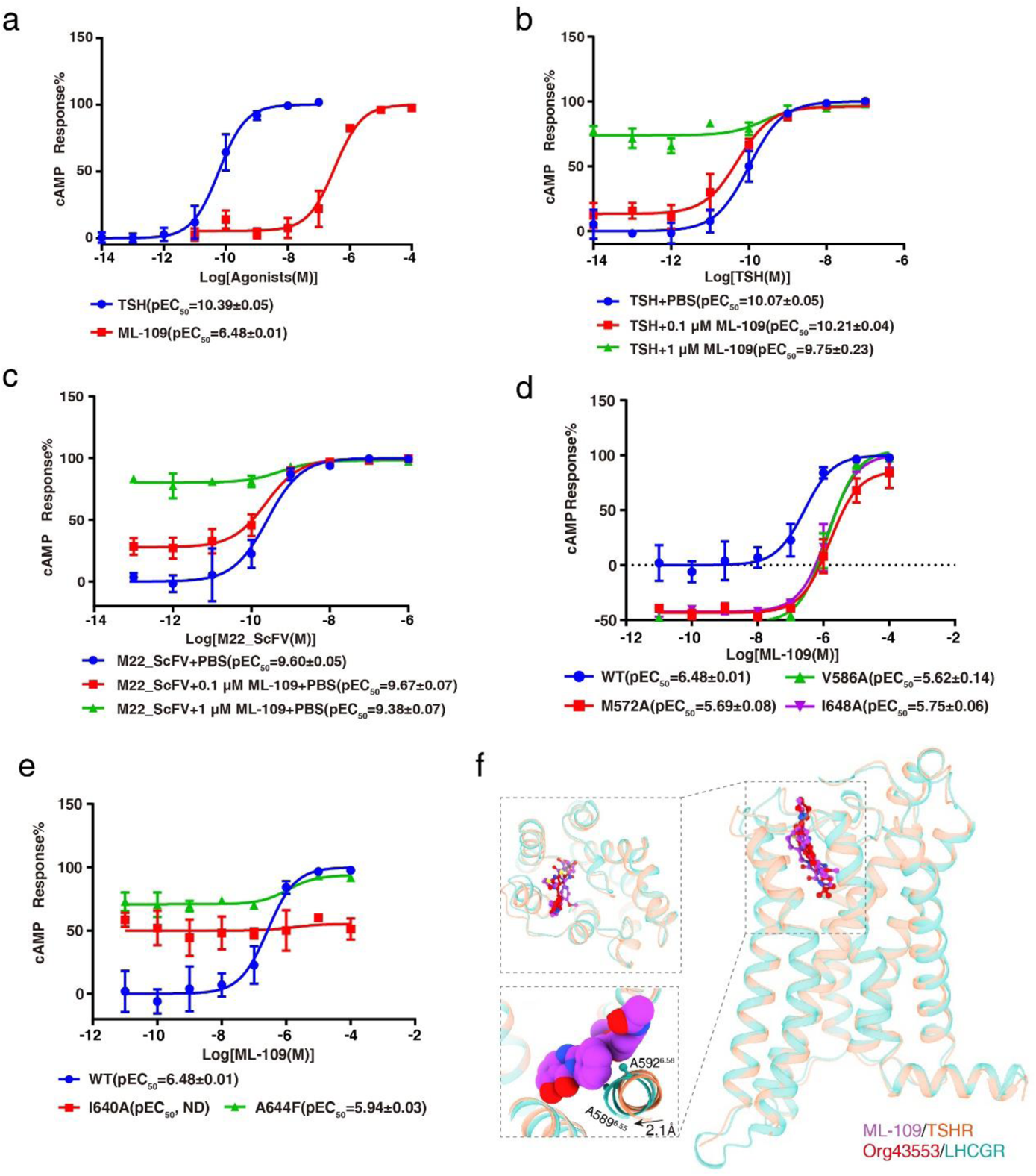
The binding pocket of ML-109 in the top half of the TMD. **a-e** Concentration-response curves for TSH, M22_ScFv and ML-109-induced TSHR activation. Data from three independent experiments are presented as the mean ± s.e.m. TSH and ML-109 induced TSHR activation, respectively (**a**); TSH plus ML-109 induced TSHR activation (**b**); M22_ScFv plus ML-109 induced TSHR activation (**c**); ML-109 alone induced mutated TSHR activation (**d-e**). **f**, Structure comparison of ML-109 binding pocket in the TSH-TSHR-Gs complex with Org43553 binding pocket in the CG-LHCGR-Gs complex.

**Extended Data Fig. 8.**
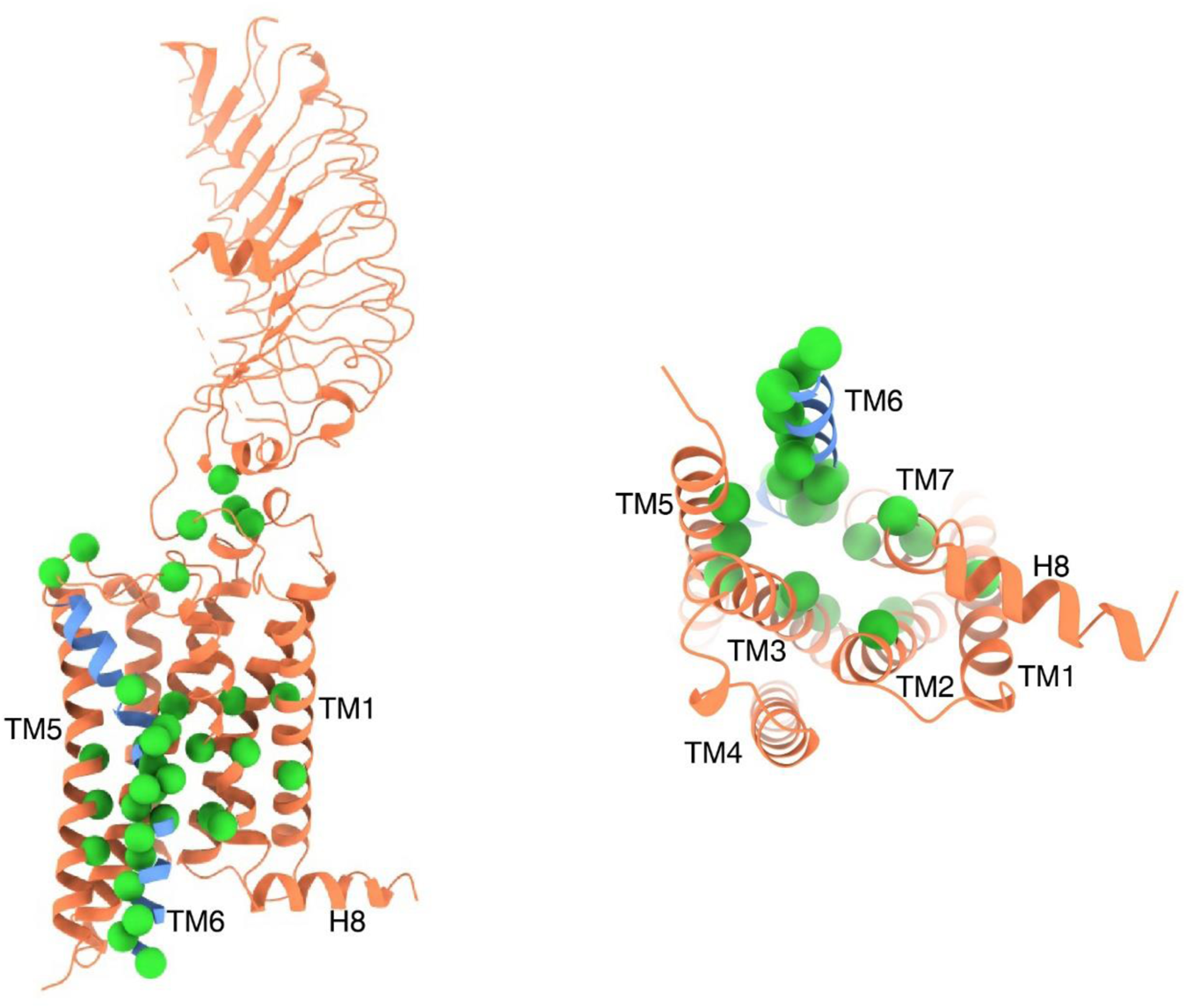
Constitutively active mutations in TSHR. The constitutively active mutations are highlighted in green spheres.

**Extended Data Table 1:**
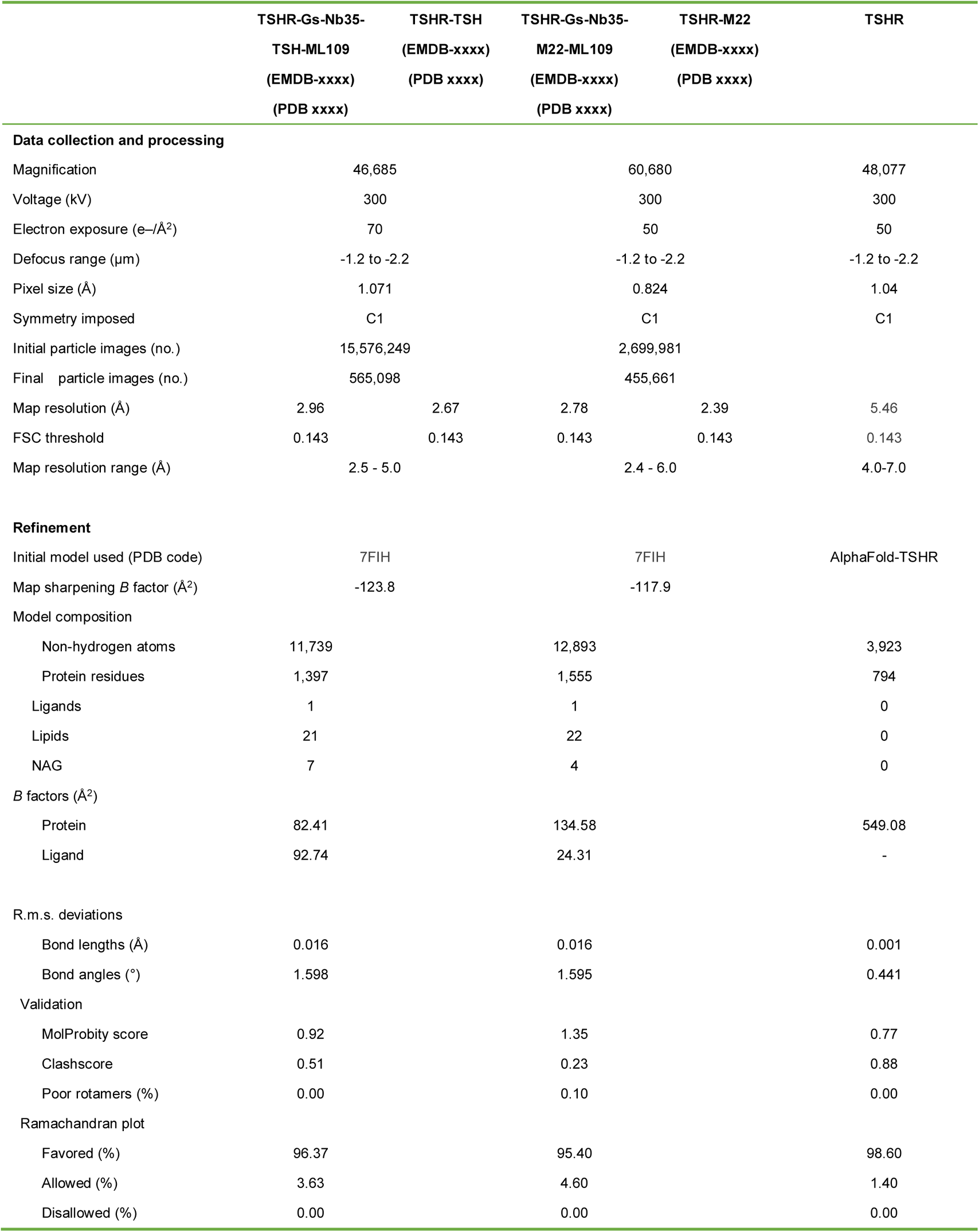
Cryo-EM data collection, refinement and validation statistics.

|  | TSHR-Gs-Nb35-TSH-ML109<br>(EMDB-xxxx)<br>(PDB xxxx) | TSHR-TSH<br>(EMDB-xxxx)<br>(PDB xxxx) | TSHR-Gs-Nb35-M22-ML109<br>(EMDB-xxxx)<br>(PDB xxxx) | TSHR-M22<br>(EMDB-xxxx)<br>(PDB xxxx) | TSHR |
| --- | --- | --- | --- | --- | --- |
| **Data collection and processing** |  |  |  |  |  |
| Magnification | 46,685 |  | 60,680 |  | 48,077 |
| Voltage (kV) | 300 |  | 300 |  | 300 |
| Electron exposure (e-/Å<sup>2</sup>) | 70 |  | 50 |  | 50 |
| Defocus range (μm) | -1.2 to -2.2 |  | -1.2 to -2.2 |  | -1.2 to -2.2 |
| Pixel size (Å) | 1.071 |  | 0.824 |  | 1.04 |
| Symmetry imposed | C1 |  | C1 |  | C1 |
| Initial particle images (no.) | 15,576,249 |  | 2,699,981 |  |  |
| Final particle images (no.) | 565,098 |  | 455,661 |  |  |
| Map resolution (Å) | 2.96 | 2.67 | 2.78 | 2.39 | 5.46 |
| FSC threshold | 0.143 | 0.143 | 0.143 | 0.143 | 0.143 |
| Map resolution range (Å) | 2.5 - 5.0 |  | 2.4 - 6.0 |  | 4.0-7.0 |
| **Refinement** |  |  |  |  |  |
| Initial model used (PDB code) | 7FIH |  | 7FIH |  | AlphaFold-TSHR |
| Map sharpening *B* factor (Å<sup>2</sup>) | -123.8 |  | -117.9 |  |  |
| Model composition |  |  |  |  |  |
| Non-hydrogen atoms | 11,739 |  | 12,893 |  | 3,923 |
| Protein residues | 1,397 |  | 1,555 |  | 794 |
| Ligands | 1 |  | 1 |  | 0 |
| Lipids | 21 |  | 22 |  | 0 |
| NAG | 7 |  | 4 |  | 0 |
| *B* factors (Å<sup>2</sup>) |  |  |  |  |  |
| Protein | 82.41 |  | 134.58 |  | 549.08 |
| Ligand | 92.74 |  | 24.31 |  | - |
| R.m.s. deviations |  |  |  |  |  |
| Bond lengths (Å) | 0.016 |  | 0.016 |  | 0.001 |
| Bond angles (°) | 1.598 |  | 1.595 |  | 0.441 |
| Validation |  |  |  |  |  |
| MolProbity score | 0.92 |  | 1.35 |  | 0.77 |
| Clashscore | 0.51 |  | 0.23 |  | 0.88 |
| Poor rotamers (%) | 0.00 |  | 0.10 |  | 0.00 |
| Ramachandran plot |  |  |  |  |  |
| Favored (%) | 96.37 |  | 95.40 |  | 98.60 |
| Allowed (%) | 3.63 |  | 4.60 |  | 1.40 |
| Disallowed (%) | 0.00 |  | 0.00 |  | 0.00 |

**Extended Data Table 2:** Ligands-induced activation on wild type and TSHR with site-directed mutations. Data represent mean pEC_50_ (pEC_50_ ± SEM), Emax (E_max_ ± SEM), E_min_ (E_min_ ± SEM). Experiments were performed in triplicate. Statistical differences between WT and mutations were determined by One-way ANOVA (and nonparametric). The E_min_ and E_max_ of TSHR mutants were normalized by WT receptor within each individual experiment, with the basal activity for WT as zero, while the fitted Emax of WT as 100, *P<0.05, **P<0.01, ***P<0.001 versus WT.

|  | Constructs | pEC<sub>50</sub> | E<sub>min</sub> | E<sub>max</sub> |
| --- | --- | --- | --- | --- |
| TSH | WT | 10.39±0.05 | 0 | 100 |
|  | K58E | 9.17±0.11** | 3.13±2.53 | 94.89±3.31 |
|  | K209A | 9.17±0.12** | -2.09±0.41 | 99.62±0.16 |
|  | Y385G | 9.33±0.08** | 16.80±1.26 | 102.58±1.75 |
|  | D386G | 9.96±0.04** | 7.57±2.81 | 102.94±3.39 |
|  | Y387G | 9.79±0.03** | 4.64±1.06 | 102.61±3.38 |
|  | Y385G,D386G,Y387G | 8.46±0.06** | 0.44±0.79 | 104.35±4.62 |
|  | Deletion of residues 291-394 | ND | ND | ND |
| TSH+PBS | WT | 10.07±0.05 | 0 | 100 |
| TSH+0.1µm ML-109 |  | 10.21±0.04 | 24.38±6.31 | 99.21±1.61 |
| TSH+1µm ML-109 |  | 9.75±0.23 | 85.85±6.91 | 101.37±2.38 |
| M22_ScFv | WT | 9.65±0.06 | 0 | 100 |
|  | Y385G | 9.79±0.05 | -2.47±0.75 | 101.53±1.29 |
|  | D386G | 9.54±0.02 | -5.30±7.99 | 101.47±3.32 |
|  | Y387G | 9.54±0.07 | -4.40±3.52 | 95.76±2.19 |
|  | Y385G,D386G,Y387G | 9.66±0.13 | 6.60±5.94 | 96.38±1.19 |
| M22_ScFv+PBS | WT | 9.60±0.05 | 0 | 100 |
| M22_ScFv +0.1µm ML-109 |  | 9.67±0.07 | 33.94±4.32 | 102.14±1.67 |
| M22_ScFv +1µm ML-109 |  | 9.38±0.07 | 85.74±2.79 | 101.5±1.82 |
| ML-109 | WT | 6.48±0.01 | 0 | 100 |
|  | V586A | 5.69±0.08** | -34.98±4.29 | 90.31±2.37 |
|  | M572A | 5.62±0.14** | -41.56±5.48 | 92.96±6.45 |
|  | I648A | ND | ND | ND |
|  | I640A | 5.94±0.03** | 72.30±2.60 | 98.75±4.02 |
|  | A644F | 5.75±0.06** | -36.39±3.16 | 103.47±2.57 |

## References

1. Dietrich, J. W., Landgrafe, G. & Fotiadou, E. H. TSH and Thyrotropic Agonists: Key Actors in Thyroid Homeostasis. J Thyroid Res 2012, 351864, doi:10.1155/2012/351864 (2012).

2. Tuncel, M. Thyroid Stimulating Hormone Receptor. Mol Imaging Radionucl Ther 26, 87–91, doi:10.4274/2017.26.suppl.10 (2017).

3. Allgeier, A. et al. The human thyrotropin receptor activates G-proteins Gs and Gq/11. J Biol Chem 269, 13733–13735 (1994).

4. Maenhaut, C. et al. Ontogeny, Anatomy, Metabolism and Physiology of the Thyroid in Endotext (eds K. R. Feingold et al.) (2000).

5. Fan, Q. R. & Hendrickson, W. A. Structural biology of glycoprotein hormones and their receptors. Endocrine 26, 179–188, doi:10.1385/endo:26:3:179 (2005).

6. Duan, J. et al. Structures of full-length glycoprotein hormone receptor signalling complexes. Nature, doi:10.1038/s41586-021-03924-2 (2021).

7. Jiang, X. et al. Structure of follicle-stimulating hormone in complex with the entire ectodomain of its receptor. Proc Natl Acad Sci U S A 109, 12491–12496, doi:10.1073/pnas.1206643109 (2012).

8. Morshed, S. A. & Davies, T. F. Graves’ Disease Mechanisms: The Role of Stimulating, Blocking, and Cleavage Region TSH Receptor Antibodies. Horm Metab Res 47, 727–734, doi:10.1055/s-0035-1559633 (2015).

9. Rapoport, B. & McLachlan, S. M. TSH Receptor Cleavage Into Subunits and Shedding of the A-Subunit; A Molecular and Clinical Perspective. Endocr Rev 37, 114–134, doi:10.1210/er.2015-1098 (2016).

10. Kopp, P. The TSH receptor and its role in thyroid disease. Cell Mol Life Sci 58, 1301–1322, doi:10.1007/pl00000941 (2001).

11. Dechairo, B. M. et al. Association of the TSHR gene with Graves’ disease: the first disease specific locus. Eur J Hum Genet 13, 1223–1230, doi:10.1038/sj.ejhg.5201485 (2005).

12. Taylor, P. N. et al. Global epidemiology of hyperthyroidism and hypothyroidism. Nat Rev Endocrinol 14, 301–316, doi:10.1038/nrendo.2018.18 (2018).

13. Neumann, S. & Gershengorn, M. C. Small molecule TSHR agonists and antagonists. Ann Endocrinol (Paris) 72, 74–76, doi:10.1016/j.ando.2011.03.002 (2011).

14. Neumann, S. et al. Small-molecule agonists for the thyrotropin receptor stimulate thyroid function in human thyrocytes and mice. Proc Natl Acad Sci U S A 106, 12471–12476, doi:10.1073/pnas.0904506106 (2009).

15. Chazenbalk, G. D. et al. Evidence that the thyrotropin receptor ectodomain contains not one, but two, cleavage sites. Endocrinology 138, 2893–2899, doi:10.1210/endo.138.7.5259 (1997).

16. Chen, C. R., Salazar, L. M., McLachlan, S. M. & Rapoport, B. Deleting the Redundant TSH Receptor C-Peptide Region Permits Generation of the Conformationally Intact Extracellular Domain by Insect Cells. Endocrinology 156, 2732–2738, doi:10.1210/en.2015-1154 (2015).

17. Ozcabi, B. et al. Testotoxicosis: Report of Two Cases, One with a Novel Mutation in LHCGR Gene. J Clin Res Pediatr Endocrinol 7, 242–248, doi:10.4274/jcrpe.2067 (2015).

18. Rasmussen, S. G. et al. Crystal structure of the beta2 adrenergic receptor-Gs protein complex. Nature 477, 549–555, doi:10.1038/nature10361 (2011).

19. Latif, R., Ando, T., Daniel, S. & Davies, T. F. Localization and regulation of thyrotropin receptors within lipid rafts. Endocrinology 144, 4725–4728, doi:10.1210/en.2003-0932 (2003).

20. Latif, R., Ando, T. & Davies, T. F. Lipid rafts are triage centers for multimeric and monomeric thyrotropin receptor regulation. Endocrinology 148, 3164–3175, doi:10.1210/en.2006-1580 (2007).

21. A., Q. R. F. a. W. Structural Biology of Glycoprotein Hormones and their Receptors. (2005).

22. Smits, G. et al. Glycoprotein hormone receptors: determinants in leucine-rich repeats responsible for ligand specificity. EMBO J 22, 2692–2703, doi:10.1093/emboj/cdg260 (2003).

23. Pierce, J. G. & Parsons, T. F. Glycoprotein hormones: structure and function. Annu Rev Biochem 50, 465–495, doi:10.1146/annurev.bi.50.070181.002341 (1981).

24. A. J. Lapthorn, D. C. H. s. Crystal structure of human chorionic gonadotropin. (1994).

25. Fox, K. M., Dias, J. A. & Van Roey, P. Three-dimensional structure of human follicle-stimulating hormone. Mol Endocrinol 15, 378–389, doi:10.1210/mend.15.3.0603 (2001).

26. Bruser, A. et al. The Activation Mechanism of Glycoprotein Hormone Receptors with Implications in the Cause and Therapy of Endocrine Diseases. J Biol Chem 291, 508–520, doi:10.1074/jbc.M115.701102 (2016).

27. Evans, M. et al. Monoclonal autoantibodies to the TSH receptor, one with stimulating activity and one with blocking activity, obtained from the same blood sample. Clin Endocrinol (Oxf) 73, 404–412, doi:10.1111/j.1365-2265.2010.03831.x (2010).

28. van Koppen, C. J. et al. Mechanism of action of a nanomolar potent, allosteric antagonist of the thyroid-stimulating hormone receptor. Br J Pharmacol 165, 2314–2324, doi:10.1111/j.1476-5381.2011.01709.x (2012).

29. Sanders, P. et al. Crystal structure of the TSH receptor (TSHR) bound to a blocking-type TSHR autoantibody. J Mol Endocrinol 46, 81–99, doi:10.1530/JME-10-0127 (2011).

30. Sanders, J. et al. Human monoclonal thyroid stimulating autoantibody. Lancet 362, 126–128, doi:10.1016/s0140-6736(03)13866-4 (2003).

31. Sanders, J. et al. Crystal structure of the TSH receptor in complex with a thyroid-stimulating autoantibody. Thyroid 17, 395–410, doi:10.1089/thy.2007.0034 (2007).

32. Kleinau, G. & Vassart, G. TSH Receptor Mutations and Diseases in Endotext (eds K. R. Feingold et al.) (2000).

33. Carpenter, B., Nehme, R., Warne, T., Leslie, A. G. & Tate, C. G. Structure of the adenosine A(2A) receptor bound to an engineered G protein. Nature 536, 104–107, doi:10.1038/nature18966 (2016).

34. Liang, Y. L. et al. Dominant Negative G Proteins Enhance Formation and Purification of Agonist-GPCR-G Protein Complexes for Structure Determination. ACS Pharmacol Transl Sci 1, 12–20, doi:10.1021/acsptsci.8b00017 (2018).

35. Maeda, S., Qu, Q., Robertson, M. J., Skiniotis, G. & Kobilka, B. K. Structures of the M1 and M2 muscarinic acetylcholine receptor/G-protein complexes. Science 364, 552–557, doi:10.1126/science.aaw5188 (2019).

36. Mastronarde, D. N. Automated electron microscope tomography using robust prediction of specimen movements. J Struct Biol 152, 36–51, doi:10.1016/j.jsb.2005.07.007 (2005).

37. Zheng, S. Q. et al. MotionCor2: anisotropic correction of beam-induced motion for improved cryo-electron microscopy. Nat Methods 14, 331–332, doi:10.1038/nmeth.4193 (2017).

38. Zhang, K. Gctf: Real-time CTF determination and correction. J Struct Biol 193, 1–12, doi:10.1016/j.jsb.2015.11.003 (2016).

39. Scheres, S. H. RELION: implementation of a Bayesian approach to cryo-EM structure determination. J Struct Biol 180, 519–530, doi:10.1016/j.jsb.2012.09.006 (2012).

40. Punjani, A., Rubinstein, J. L., Fleet, D. J. & Brubaker, M. A. cryoSPARC: algorithms for rapid unsupervised cryo-EM structure determination. Nat Methods 14, 290–296, doi:10.1038/nmeth.4169 (2017).

41. Pettersen, E. F. et al. UCSF Chimera--a visualization system for exploratory research and analysis. J Comput Chem 25, 1605–1612, doi:10.1002/jcc.20084 (2004).

42. Emsley, P. & Cowtan, K. Coot: model-building tools for molecular graphics. Acta Crystallogr D Biol Crystallogr 60, 2126–2132, doi:10.1107/S0907444904019158 (2004).

43. Croll, T. I. ISOLDE: a physically realistic environment for model building into low-resolution electron-density maps. Acta Crystallogr D Struct Biol 74, 519–530, doi:10.1107/S2059798318002425 (2018).

44. Adams, P. D. et al. PHENIX: a comprehensive Python-based system for macromolecular structure solution. Acta Crystallogr D Biol Crystallogr 66, 213–221, doi:10.1107/S0907444909052925 (2010).

45. Chen, V. B. et al. MolProbity: all-atom structure validation for macromolecular crystallography. Acta Crystallogr D Biol Crystallogr 66, 12–21, doi:10.1107/S0907444909042073 (2010).

46. Pettersen, E. F. et al. UCSF ChimeraX: Structure visualization for researchers, educators, and developers. Protein Sci, doi:10.1002/pro.3943 (2020).

47. Lomize, M. A., Pogozheva, I. D., Joo, H., Mosberg, H. I. & Lomize, A. L. OPM database and PPM web server: resources for positioning of proteins in membranes. Nucleic Acids Res 40, D370–376, doi:10.1093/nar/gkr703 (2012).

48. Jo, S. et al. CHARMM-GUI 10 years for biomolecular modeling and simulation. J Comput Chem 38, 1114–1124, doi:10.1002/jcc.24660 (2017).

